# Defective *Slc7a7* transport reduces erythropoietin compromising erythropoiesis and iron homeostasis

**DOI:** 10.1101/2021.08.15.456393

**Authors:** Judith Giroud-Gerbetant, Fernando Sotillo, Gonzalo Hernández, Cian J Lynch, Irene Ruano, Barbara Siri, David Sebastian, Antonio Zorzano, Rafael Artuch, Aida Ormazabal, Mayka Sánchez, Günter Weiss, Neus Prats, Carlo Dionisi-Vici, Manuel Serrano, Manuel Palacín, Susanna Bodoy

**Author notes:** Susanna Bodoy, Institute for Research in Biomedicine Barcelona, Baldiri Reixac, 12, Barcelona 08028, Spain; Manuel Palacín, Institute for Research in Biomedicine Barcelona, Baldiri Reixac, 12, Barcelona 08028, Spain. **Author Contributions:** J.G., F.S. and S.B. designed and performed experiments, and interpreted and analyzed data. J.G., S.B., and M.P. designed the research and wrote the manuscript, with input from all the authors. J.C., C.L., I.R., B.S., D.S., A.O. performed research. G.W., A.Z., R.A., M.S., N.P., C.D.V., and M.S. provided reagents and provided intellectual input. **Competing Interest Statement:** The authors declare that they have no conflict of interest.

## Abstract

Lysinuric Protein Intolerance (LPI) is an inborn error of metabolism resulting from *SLC7A7* deficiency that causes diminished plasma concentration of cationic amino acids. The clinical picture is highly heterogeneous among patients, who commonly present intolerance to protein intake and more severe complications such as hematological abnormalities and kidney failure. Although current treatments aim to address the metabolic defects of LPI, they have been unsatisfactory when treating the most severe symptoms. Here we show that the absence of *Slc7a7* in mice causes iron overload as a result of erythropoiesis failure. Regarding iron metabolism, we demonstrate that reduced plasma erythropoietin triggers a strong iron overload, as erythropoietin administration restores normal iron levels and mitigate hematological alterations. Interestingly, we found that human LPI is associated with hyperferritinemia but not iron overload, a trait that might be influenced by the citrulline treatment. Furthermore, we show that erythropoietin is a key factor in the hematological abnormalities in LPI. Our study reveals a mechanism leading to LPI-induced hematological complications and identifies erythropoietin supplementation as a promising therapeutic strategy for human LPI.

**Significance Statement:** The systemic metabolic environment derived from *Slc7a7*-ablation in epithelial cells from kidney and intestine causes erythropoiesis failure prompting therefore iron overload. Here, we identify erythropoietin as the main driver of erythropoiesis failure as exogenous erythropoietin administration restores normal erythroblast population. In addition, we have also analyzed human data and found that patients with LPI have abnormal ferritin levels. Finally, as human LPI, citrulline treatment in mice restores normal iron homeostasis, highlighting the relevance of the systemic environment in LPI. Erythropoietin supplementation emerges as a promising therapeutic strategy for human LPI without the inflammatory effect associated with citrulline supplementation.

## Introduction

Amino acids are key precursors for the synthesis of proteins and other important compounds, such as peptide hormones and neurotransmitters. However, an imbalance in amino acid intake may compromise metabolic processes involving polyamines, the urea cycle and therefore nitrogen recycling (1). Here, the kidney plays an important role by participating in the reabsorption of amino acids and the exchange of nitrogenous metabolites. As such, amino acid transporters are membrane-bound proteins that orchestrate the transfer of amino acids in and out of the cell, and mutations in human amino acid transporters are associated with severe metabolic disorders (2).

*SLC7A7* encodes for y^+^LAT1, a light subunit of the heteromeric amino acid transporter family. y^+^LAT1 mediates the exchange of cationic amino acids with neutral amino acids plus sodium across the basolateral membrane of epithelial cells. Loss-of-function in *SLC7A7* causes lysinuric protein intolerance (LPI, MIM 222700), a rare autosomal recessive disease characterized by a pleiotropic phenotype (3). Patients with LPI’s symptoms vary from vomiting, failure to thrive, and hepatosplenomegaly to more severe symptoms, including lung complications, hematological abnormalities, and kidney failure. To date, the standard treatment for LPI is a low protein-based diet supplemented with oral citrulline, which is converted to arginine in renal epithelial cells, thereby correcting hypoargininemia, defects in the urea cycle, and hyperammonemia (4).

The metabolic alterations derived from LPI are well-known and have been extensively studied (5). However, the implications of the systemic metabolic environment resulting from SLC7A7 mutations (i.e., hypoargininemia and hyperammonemia) in the plethora of hematological and immune abnormalities in patients with LPI remains poorly understood. Among these hematological anomalies, cytopenia and normochromic or hypochromic-microcytic anemia (6) are commonly found in patients with LPI. However, these individuals also frequently present increased levels of ferritin (4). While such conditions are not life-threatening when controlled, they become harmful when the disease progresses. An increasing body of literature has shown how iron metabolism is tightly linked to red blood cell (RBC) maturation and recycling. First, during the early stage of erythroblast maturation, proerythroblasts start producing hemoglobin. Then, throughout the proliferation and differentiation process, macrophages provide iron to the erythroblasts (7) to finally produce reticulocytes, which will finally mature in the bloodstream.

Contrary, upon RBC aging, resident macrophages in the spleen take up erythrocytes to further trigger hemoglobin breakdown and, ultimately, iron recycling and release (8). Various immunological and hematological mechanisms are activated during anemia to fulfill erythropoiesis, and, in turn, they contribute to increased RBC production. Among these phenotypic changes, the fine-tuning of erythropoietin production is one of the earliest events to occur (9).

Erythropoietin is a glycoprotein that is synthesized mainly in the kidney. This process is regulated by two important factors. First, erythropoietin synthesis is oxygen-dependent.

Specifically, hypoxia-inducible factor 2 (HIF-2) stabilization activates erythropoietin transcription. Second, erythropoietin production is also directly linked to the number of erythropoietin-producing cells (REPCs) in the kidney (10). This extremely rare type of cell is localized within the peritubular interstitial space and is known to express the PDGFR-ß polypeptide and CD73 established pericyte markers (11). In disease conditions, such as chronic kidney disease (CKD) and/or end-stage renal disease, erythropoietin levels are decreased and are consequently unable to meet the demands for erythropoietin, thereby perpetuating anemia. The end-stage of many renal pathologies is renal fibrosis, which has been proposed as a plausible mechanism for decreased erythropoietin production in CKD and kidney injury (12).

Using a genetically modified mouse model of LPI (13), we found that erythropoietin deficiency is at the root cause of the hematological abnormalities experienced by patients of LPI. Our analysis of the LPI mouse model (*Slc7a7^-/-^* mice) demonstrated that after *Slc7a7* ablation mice developed defective erythropoiesis and iron overload. However, ablation of *Slc7a7* specifically in the myeloid-cell lineage revealed that this phenotype was not caused by a cell-autonomous defect of *Slc7a7* expression in macrophages. Interestingly, our findings suggests that erythropoietin is the underlying of iron accumulation in mice, as erythropoietin secretion induced by exogenous erythropoietin administration and phenylhydrazine restored a normal iron phenotype and erythropoiesis. Moreover, in patients with LPI, we found that RBC abnormalities were highly heterogeneous, as 50% of them showed increased RBC counts whereas the other 50% presented anemia, as determined by decreased hemoglobin content and total hematocrit.

As 90% of the patients were on citrulline therapy, we treated *Slc7a7^-/-^* mice with citrulline supplementation. This approach restored the iron phenotype, thereby further suggesting that the differences observed in the untreated *Slc7a7^-/-^* mice and the patients were probably because of citrulline therapy. In sum, this study provides evidence of an iron related phenotype in *Slc7a7^-/-^* mice that is probably masked in patients with LPI due to citrulline therapy and suggesting that erythropoietin supplementation could be a concomitant therapy for the hematological abnormalities of human LPI.

## Results

### *Slc7a7^-/-^* mice recapitulate LPI-associated hematological alterations and develop iron overload

We previously generated *a Slc7a7^-/-^* mouse model in which Cre recombinase is ubiquitously expressed after tamoxifen administration (13)(Fig 1A) and showed that these mice model LPI, a rare autosomal disorder characterized by a multifaceted immunological and hematological defect.

**Figure 1.**
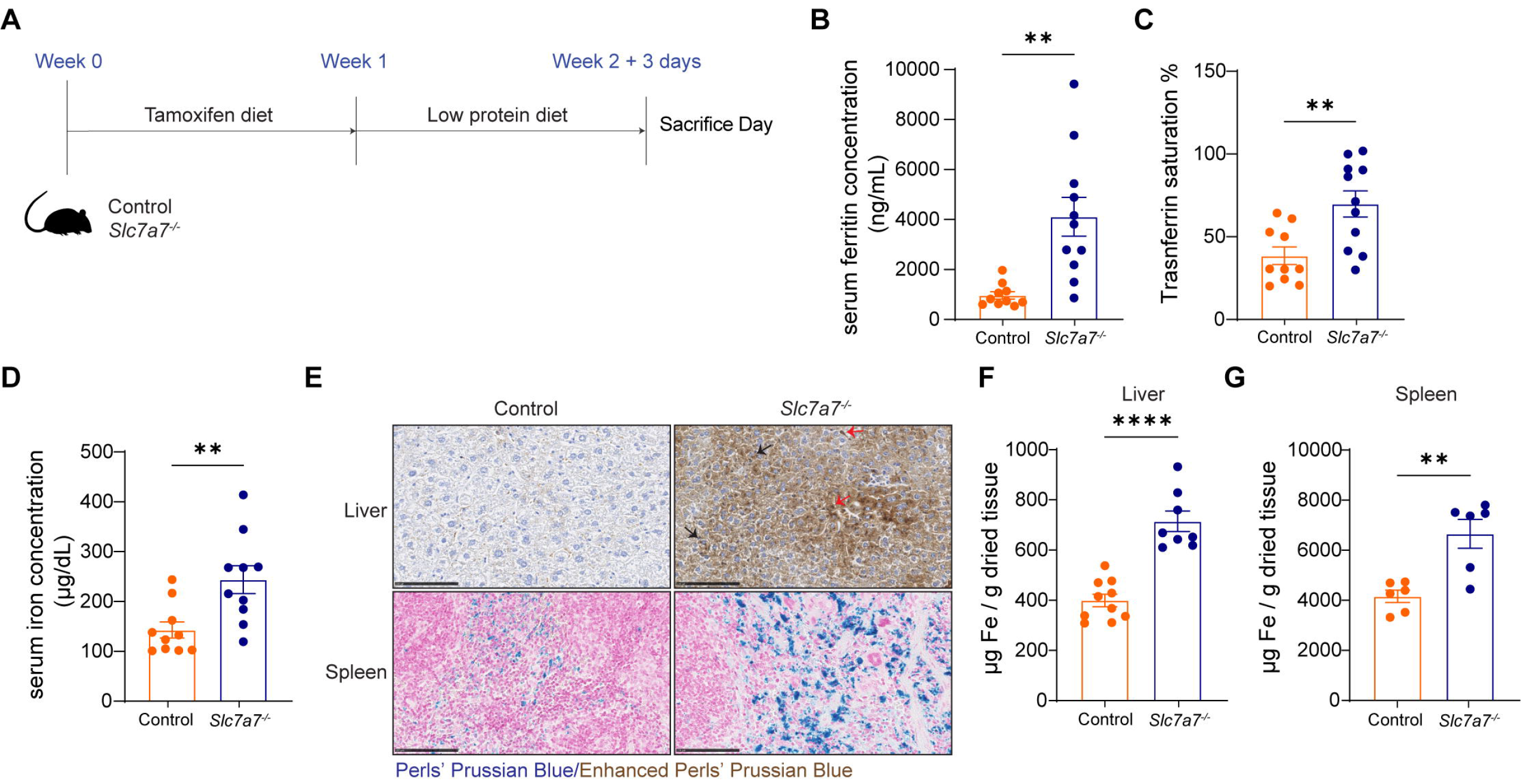
***Slc7a7* knockout mice develop iron overload. (**A) Graphical scheme of tamoxifen-inducible *Slc7a7* mouse model. (B) Enzyme-linked immunosorbent assay (ELISA) of serum ferritin in controls and *Slc7a7^-/-^* mice. (C) Serum transferrin saturation in *Slc7a7^-/-^* mice and control littermates. (D) Serum iron concentration in control and *Slc7a7^-/-^* mice. (E) Upper panel: Enhanced Perls Prussian blue staining in liver sections. Brown indicates iron staining in hepatocytes (black arrow) and Kupffer cells (red arrow). Lower panel: Perls Prussian blue staining in spleen sections. Hematoxylin and eosin staining was used as a background. Scale (F-G) Spectrophotometric analysis of total iron content in liver and spleen from bars: 100 μ control and *Slc7a7 -/-* mice, respectively. Data information: Data are mean ± SEM. **p < 0.01, ****p < 0.0001 (two-tailed unpaired student-t test). Each data point represents a single animal.

Regarding the hematological complications, patients with LPI develop normochromic and/or hypochromic/microcytic anemia (6). Here we first tested whether the inducible *Slc7a7^-/-^* mice also develop similar abnormalities. Whole blood count analysis revealed that mean corpuscular volume (MCV) and mean corpuscular hemoglobin (MCH) were significantly lower in *Slc7a7^-/-^* mice than in controls, but no differences were evident in hemoglobin concentration, RBC count, or hematocrit (HCT) (Table 1). Similarly, histological analysis of peripheral blood showed that erythrocytes from *Slc7a7^-/-^* mice were significantly smaller (Fig S1A). Some patients with LPI also have elevated ferritin levels in serum (14). Accordingly, *Slc7a7^-/-^* mice had significantly higher amounts of serum ferritin (Fig 1B). Serum ferritin is a well-known marker of inflammatory disease (15). To evaluate whether greater ferritin levels were paralleled by an increased inflammatory phenotype, we examined changes in inflammatory cytokines in serum from control and *Slc7a7^-/-^* mice. We did not detect obvious changes between the two genotypes (Fig S1B). Consistently, we did not observe significant changes in the gene expression of key interleukins and cytokines in sorted red pulp macrophages (RPMs) from control and *Slc7a7^-/-^* mice, except *Ccl5* which showed decreased gene expression levels compared to controls (Fig S1C). Noteworthy, decreased *Ccl5* expression is a trait also found in patients with acquired aplastic anemia (16).

**Table 1.**
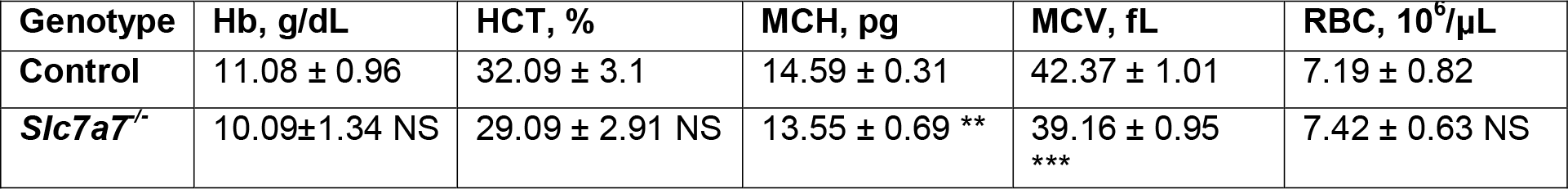
Whole blood count analysis in *Slc7a7-/-* mice. Hemoglobin (Hb), hematocrit (Hct), mean corpuscular hemoglobin (MCH), mean corpuscular volume (MCV), and red blood cell counts (RBC) from control and *Slc7a7-/-* mice. Data are ± SEM. ** P ≤ 0.01, *** P ≤ 0.001 between genotypes and NS stands for non-significant. P-val were calculated using a two-tailed Student’s ttest.

Serum ferritin concentration alongside transferrin saturation in serum can be used to determine the amount of iron stored in the body. Therefore, we measured transferrin saturation in serum, which is used to determine the amount of serum iron bound to transferrin (17), and observed that transferrin saturation was significantly higher in *Slc7a7^-/-^* mice compared to control animals (Fig 1C). We next measured the serum iron levels. Like ferritin and transferrin saturation, we found that serum iron concentrations were significantly higher in *Slc7a7^-/-^* mice (Fig 1D). In addition to increased levels of plasma ferritin, and iron and transferrin saturation in serum, iron overload is also associated with aberrant iron deposits in various organs such as the liver (18). Histological analysis with enhanced Perls’ Prussian blue, which detects iron (19), showed that the hepatocytes of *Slc7a7^-/-^* mice had significantly higher levels of iron (Fig 1E and S1D). Iron deposits could also be observed in the spleen (Fig 1E and S1D). As expected, iron deposits were not detected in control animals. We next used a spectrophotometric assay to quantify iron levels in the liver and spleen and found that *Slc7a7^-/^*^-^ mice had elevated iron levels compared to control animals (Figs 1F-G).

Overall, these results indicate that *Slc7a7^-/-^* mice recapitulates the hematological alterations found in patients with LPI, associated to iron overload in different tissues and hyperferritinemia.

### Decreased FPN1 expression drives iron accumulation in red pulp macrophages

As *Slc7a7* is expressed in macrophages (20) and RPMs are the cells in charge of iron recycling in the spleen, we initially suspected that iron accumulation might be directly linked to these cells (21). Whereas enhanced Perls’ Prussian blue from liver sections demonstrated iron accumulation in hepatocytes and Kupffer cells (Fig 1C), co-staining of the macrophage marker F4/80 and Perls’ Prussian blue in spleen sections proved that in the spleen tissue iron was mainly located in the cytoplasm of the splenic RPMs (Fig 2A).

**Figure 2.**
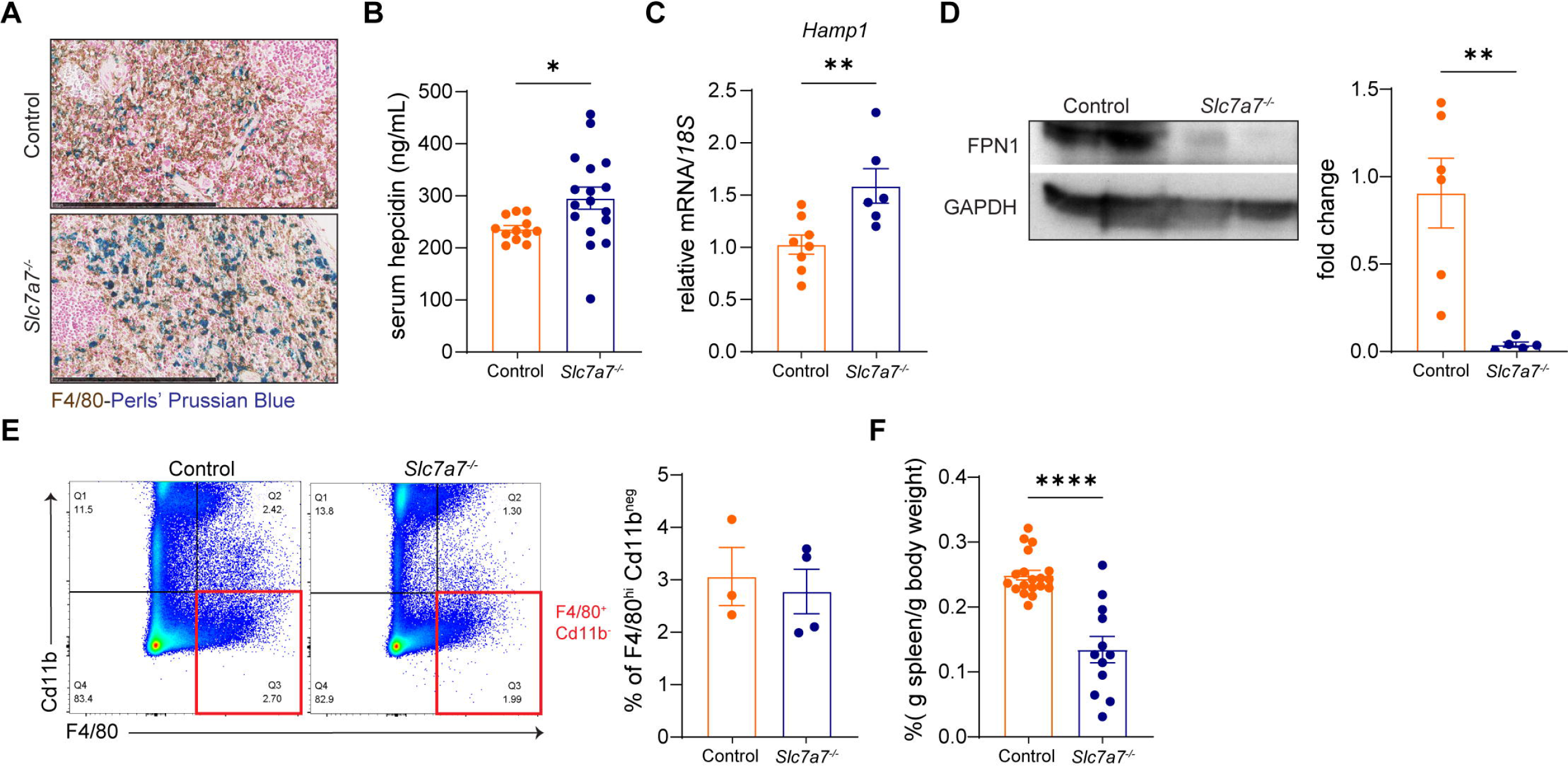
*Slc7a7* knockout mice develop iron accumulation in splenic macrophages. (A) Co-staining of the macrophage marker F4/80(brown) and Perls Prussian blue in spleen sections. Hematoxylin and eosin staining was used as a background. Scale bars: 250 μm. (B) ELISA of serum hepcidin in control and *Slc7a7^-/-^* mice. (C) Real-time RT-PCR analysis of *Hamp1* expression in liver. (D) Left panel: Western blot analysis of Fpn-1 in total homogenate from spleen tissue. Right panel: FPN-1 quantification. (E) Flow cytometry analysis of red pulp macrophages (F4/80^+^Cd11b^-^). Window in Q3 represents the red pulp macrophages population. (D) Relative spleen weight of control and *Slc7a7^-/-^* mice. Data information: Data are mean ± SEM. *p < 0.05, **p < 0.01, ***p < 0.001, ****p < 0.0001 (two-tailed unpaired student-t test). Each data point represents a single animal.

Iron metabolism in mammals is a tightly regulated process where several proteins are key for proper iron homeostasis. Ferroportin-1 is the only known iron exporter and it is post-translationally regulated by the hormone hepcidin (22). We found that serum hepcidin protein concentration was significantly higher in *Slc7a7^-/-^* mice compared to control animals (Fig 2B). In addition, hepatic hepcidin transcript (*Hamp1*) levels were also significantly increased in *Slc7a7^-/-^* mice (Fig 2C). We next measured FPN1 protein levels in *Slc7a7^-/-^* and control mice. Accordingly, western blot analysis revealed a massive decrease of FPN1 in total spleen homogenate (Fig 2D). Next, we tested whether the dramatic decrease in FPN1 protein levels was associated with a reduced RPM population (F4/80^+^Cd11b^-^). *Slc7a7^-/-^* mice had normal relative RPM numbers (Fig 2E). However absolute counts of RPMs were significantly diminished in *Slc7a7^-/-^* mice (Figs S2A- B). Histological analysis confirmed similar levels of F4/80^+^ cells in spleen sections (Fig S2C). One possibility that could account for the decreased absolute RPM cell type in *Slc7a7^-/-^* mice is reduced spleen weight. In this regard, *Slc7a7^-^*^/-^ mice showed a dramatic reduction in relative and absolute spleen size and weight (Fig 2F and S2D), yet no changes were observed for the body weight ratio of the gastrocnemius, kidney, and liver (Figs S2E-G).

### We next sought to elucidate the mechanisms underlying iron overload in *Slc7a7^-/-^* mice

To this end, we compared the transcriptome profile of sorted RPMs from control and *Slc7a7^-/-^* mice. Analysis revealed that the RPMs from both genotypes express similar levels of key- associated RPM genes (21) (Fig S2H-I). Given that RPMs are also in charge of removing senescent erythrocytes by phagocytosis and heme recycling, we compared the heme transcriptional signature (Hallmark Heme Metabolism) in sorted RPMs from control and *Slc7a7^-/-^* animals using gene set enrichment analysis (GSEA). Sorted RPMs from *Slc7a7^-/-^* mice showed a tendency of downregulation of heme signature genes, yet the overall transcriptional signature was similar between the two genotypes (Fig S2J). Taken together, these data suggest that the decreased FPN1 expression drives acute iron accumulation in RPMs.

### Partial reduction of *Slc7a7* in splenic macrophages does not cause iron overload

How does *Slc7a7* expression in the myeloid cell lineage influence iron overload? *Slc7a7* is expressed in macrophages, and the similar transcriptome profile of sorted RPMs from control and *Slc7a7^-/-^* mice suggested that iron overload is not caused by impaired iron metabolism or by changes in RPMs. We hypothesized that the systemic environment (i.e., reduced CAA in plasma and hyperammonemia) caused by *Slc7a7* deficiency in epithelial cells from the intestine and kidney orchestrates iron overload. To test this notion, we generated a myeloid-conditional *Slc7a7* knockout mouse in which Cre expression was under the control of the lysozyme 2 endogenous promoter (Lyz2Cre/Cre mice) (hereafter referred to as the *Slc7a7^LysM-/-^*). We found that *Slc7a7* expression was fully ablated in alveolar macrophages and bone marrow-derived macrophages (BMDMs). However, *Slc7a7* ablation in RPMs from *Slc7a7^LysM-/-^* was only partial, reaching similar levels of ablation to those in global *Slc7a7^-/-^* mice (Fig S3A). As expected, plasma concentration of arginine and orotic acid were similar between control and *Slc7a7^LysM-/-^* mice (Figs S3B-C) monitoring normal amino acid re-absorption and the urea cycle. We then found that myeloid-specific *Slc7a7^LysM-/-^* mice displayed similar iron levels in spleen and liver sections to those of wild-type controls (Fig S3D). Accordingly, control and *Slc7a7^LysM-/-^* mice showed similar levels of liver and spleen iron content (Fig S3E-F). Serum iron and ferritin, as well as transferrin saturation levels, were all unchanged in *Slc7a7^LysM-/-^* mice (Figs S3G-I). The results from iron levels in serum and tissue suggested that *Slc7a7* expression in macrophages is unlikely to be the main factor driving iron overload. However, we cannot rule out that the phenotype could be exacerbated with complete *Slc7a7* ablation in RPMs from the macrophage-specific cell line.

### *Slc7a7* knockout mice show failed erythropoiesis

Iron overload has been related to ineffective erythropoiesis, which is the imbalance between erythroid proliferation and differentiation. During stress erythropoiesis, the kidney hormone erythropoietin is upregulated to meet the demands of increased RBC expansion (23). We have previously shown that erythrocytes from *Slc7a7^-/-^* mice have RBC abnormalities as depicted by reduced MCV and MCH (Table 1); however, no anemia was fully evident two weeks after *Slc7a7* ablation. We hypothesized that global *Slc7a7* ablation causes ineffective erythropoiesis, which resulted in erythrocytes with a reduced MCV and MCH, which ultimately trigger iron overload. Although *Slc7a7^-/-^* mice showed normal RBC count (Table 1), they had a significant decrease in reticulocyte percentage in blood (Fig 3A).

**Figure 3.**
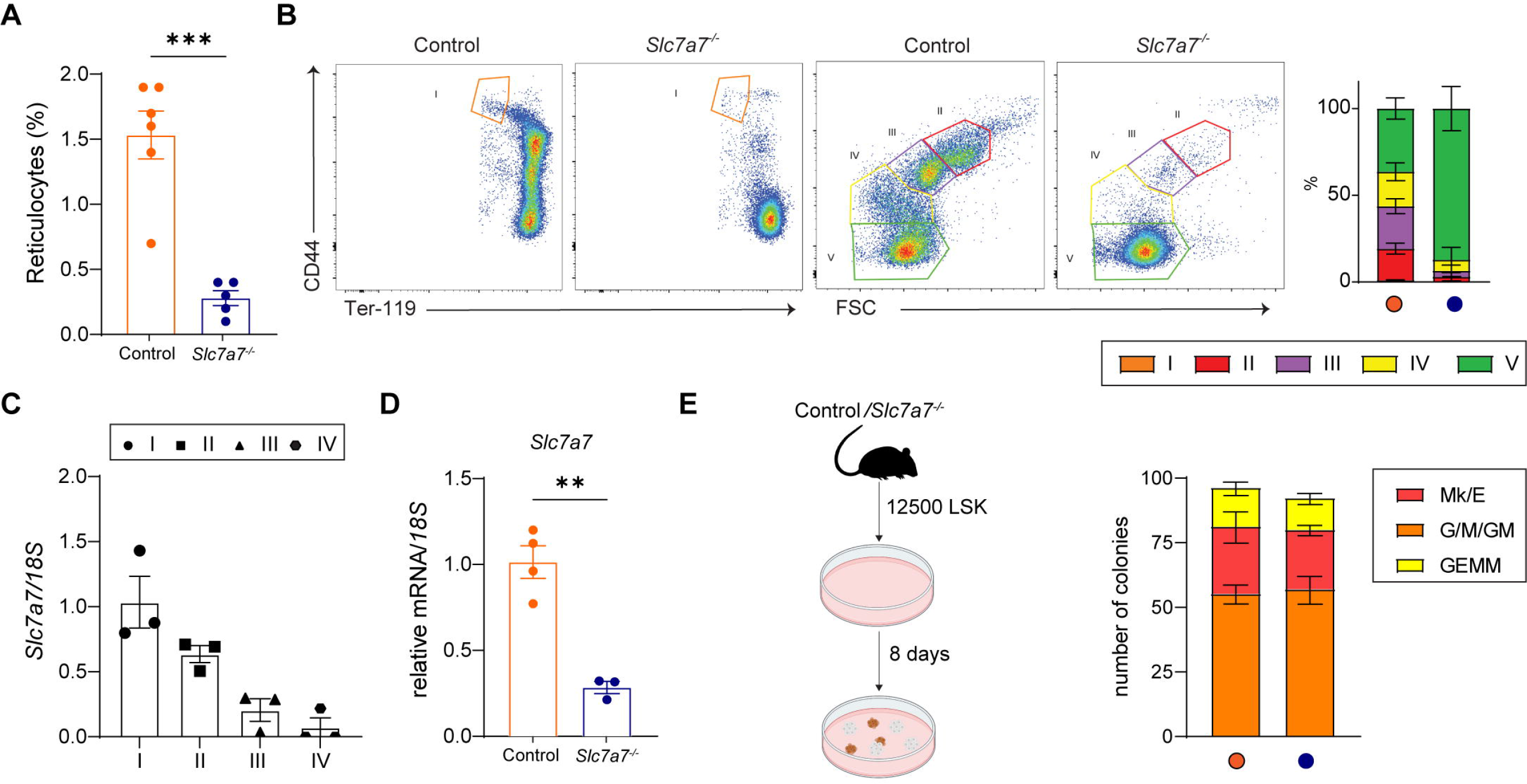
***Slc7a7* knockout mice have deficient erythropoiesis**. (A) Reticulocyte count in whole blood (as a number of reticulocytes per 1000 RBCs). (B) Left: Representative dot plots show the gating strategy for erythroid progenitors (V (*p*-val < 0,0001), IV (*p*-val < 0,0001), III (*p*- val < 0,0001), II (*p*-val < 0,01) and I (*p*-val < 0,9)) (Chen) from the indicated genotype. Briefly, cells were first gated in Ter-119^+^ and further separated by CD44 vs. Forward Scatter (FSC-A). Right: Percentage of the cell populations analyzed, n = 5. (C) Real-time PCR analysis of *Slc7a7* in sorted erythroid progenitors from control mice (IV, III, II and I). (D) Real-time PCR analysis of *Slc7a7* in sorted erythroid progenitors from control and *Slc7a7^-/-^* mice. (E) 12500 LSK cells from control and *Slc7a7^-/-^* mice were plated in Methocult for colony-forming unit (CFU). The analysis was done 8 days after the plating. G/M/GM, G granulocyte, M monocyte, GM granulocyte-monocyte; Mk/E, Mk megakaryocyte, E erythroid; and GEMM, granulocyte-erythroid-monocyte-megakaryocyte mixed CFU (n=4). Data information: Data are mean ± SEM. **p < 0.01, ***p < 0.001 (two-tailed unpaired student-t test). Each data point represents a single animal.

We next analyzed erythropoietic precursors in the bone marrow (BM) of control and *Slc7a7^-/-^* mice by flow cytometry. Of note, we found that the erythroblast number in BM was severely reduced in *Slc7a7^-/-^* mice. Specifically, these animals showed a considerably lower number of basophilic erythroblasts (region II), polychromatic erythroblasts (region III) and orthochromatic erythroblasts (region IV) than control mice, whereas the percentage of mature RBCs (region V) was significantly higher and the percentage of proerythroblasts (region I) was similar between genotypes (Fig 3B). To test whether *Slc7a7^-/-^* mice showed a different expression pattern of CD44 and CD71, we analyzed the expression of the transferrin receptor CD71 in erythropoietin progenitors. We found that the number of erythropoietic precursors at stages (II-IV) was substantially reduced in *Slc7a7^-/-^* mice (Fig S4A). These findings indicate that, despite having fewer erythrocyte precursors, *Slc7a7^-/-^* mice and control mice have the same number of mature RBCs in the blood (Table 1). Taken together, these data suggest that global ablation of *Slc7a7* causes inefficient erythropoiesis, as revealed by fewer erythroblast precursors and reticulocytopenia. In the event of ineffective BM erythropoiesis, alternative erythropoiesis is required, and extramedullary stress erythropoiesis (EMH) takes place in the spleen (23).

Therefore, we measured stress erythropoiesis in the spleen. Wild-type and *Slc7a7^-/-^* mice showed similar levels of erythroid progenitors (Fig S4B). Histological analysis confirmed that control and knockout mice presented minimal levels of EMH in splenic red pulp. However, we did detect a decrease in the size and cellularity of the red pulp (Fig S4C). Taken together, similar levels of EMH and reduced size and weight of the spleen suggested that it is unlikely that the spleen could take over upon ineffective erythropoiesis in BM.

Tusi et al., using single-cell RNA sequencing, reported how *Slc7a7* expression decreased upon erythroblast maturation (24). We therefore studied *Slc7a7* transcript during erythroblast maturation and confirmed that proerythroblasts (I) have higher *Slc7a7* expression than orthochromatic erythroblasts (IV) (Fig 3C). We then examined whether *Slc7a7* was efficiently deleted in whole erythroblast population in the BM of *Slc7a7^-/-^* mice. Gene expression analysis revealed that upon tamoxifen administration, *Slc7a7* expression in erythroblasts of *Slc7a7^-/-^* mice decreased by about 80% (Fig 3D). Next, we used in vitro colony-forming unit (CFU) assay to test the potential of sorted Lin^-^Sca1^+^cKit^+^ (LSK) cells from control and *Slc7a7^-/-^* mice to proliferate and differentiate. No differences were observed in the type or the total number of colonies between the two genotypes (Fig 3E). These results exclude a cell-autonomous basis for the deficient erythropoiesis in *Slc7a7^-/-^* mice.

### *Slc7a7^-/-^* mice have reduced erythropoietin plasma levels

Erythroferrone (ERFE) is a hormone produced by erythroid progenitors and spleen during enhanced erythropoiesis (25). ERFE downregulates liver hepcidin (*Hamp*) to increase iron availability. Interestingly, we observed that *ERFE* transcript levels in sorted erythroid progenitors were significantly lower in *Slc7a7^-/-^* mice (Fig S5A). However, plasma ERFE levels were undetectable in both control and *Slc7a7^-/-^* animals (Fig 4A), as ERFE production increases only under circumstances of enhanced erythropoiesis or erythropoietin stimulation (25). Given that erythropoietin is the hormone responsible for promoting the survival, proliferation, and differentiation of immature RBCs (26), we measured the levels of circulating erythropoietin and total erythropoietin in kidney extracts from animals to determine whether the smaller erythroid progenitor population in *Slc7a7^-/-^* mice was associated with lower serum levels. Serum and total kidney erythropoietin levels were significantly lower in *Slc7a7^-/-^* mice than in wild-type mice (Figs 4B-C). Reduced kidney erythropoietin levels in *Slc7a7^-/-^* mice were confirmed by gene expression analysis (Fig S5B). We next measured erythropoietin gene expression levels in liver and found that the erythropoietin transcript was undetectable in both genotypes (Fig S5C). This observation is consistent with the fact that liver erythropoietin levels can be detected only in embryonic stages and/or in anemic mice (27). In addition to erythropoietin, Insulin-like growth factor 1 (IGF-1) is also required for optimal erythropoiesis (28). IGF-1 stimulates erythropoiesis, thereby promoting erythroblast proliferation, and importantly, IGF-1 is required for the end-terminal enucleation of erythroid precursors. Analysis of IGF-1 levels in plasma from *Slc7a7^-/-^* and control mice revealed a higher concentration in *Slc7a7^-/-^* (Fig 4D), in contrast with what was previously reported, which could be explained by the differences in the mouse model used (29) . To rule out a possible cell-intrinsic effect of *Slc7a7* expression in the erythroblast lineage we generated *Er*GFP*creSlc7a7^LoxP/LoxP^* mice (referred to herein as *Slc7a7^EpoR-/-^*), in which the *Slc7a7* locus is specifically ablated in the erythroid lineage (Heinrich et al., 2004), by crossing *Er*GFP*cre* and *Slc7a7^flox^*^/*flox*^ mice (Fig S5D). We found that *Slc7a7* expression was reduced by about 90% in sorted erythroblasts (Ter-119^+^ cells) from *Slc7a7EpoR^-/-^* mice (Fig S5E); however, erythropoiesis deficiency was not present in the *Slc7a7EpoR^-/-^* mice (Fig S5F). In addition, whole blood count analysis showed that *Slc7a7EpoR*^-/-^ mice had normal HCT, RBC numbers, and MCV and MCH (Fig S5G-J), thereby indicating that the impaired erythropoiesis in *Slc7a7^-/-^* mice was directly linked to the systemic environment of global *Slc7a7* ablation rather than a cell-intrinsic effect of *Slc7a7* in the erythroblastic lineage.

**Figure 4.**
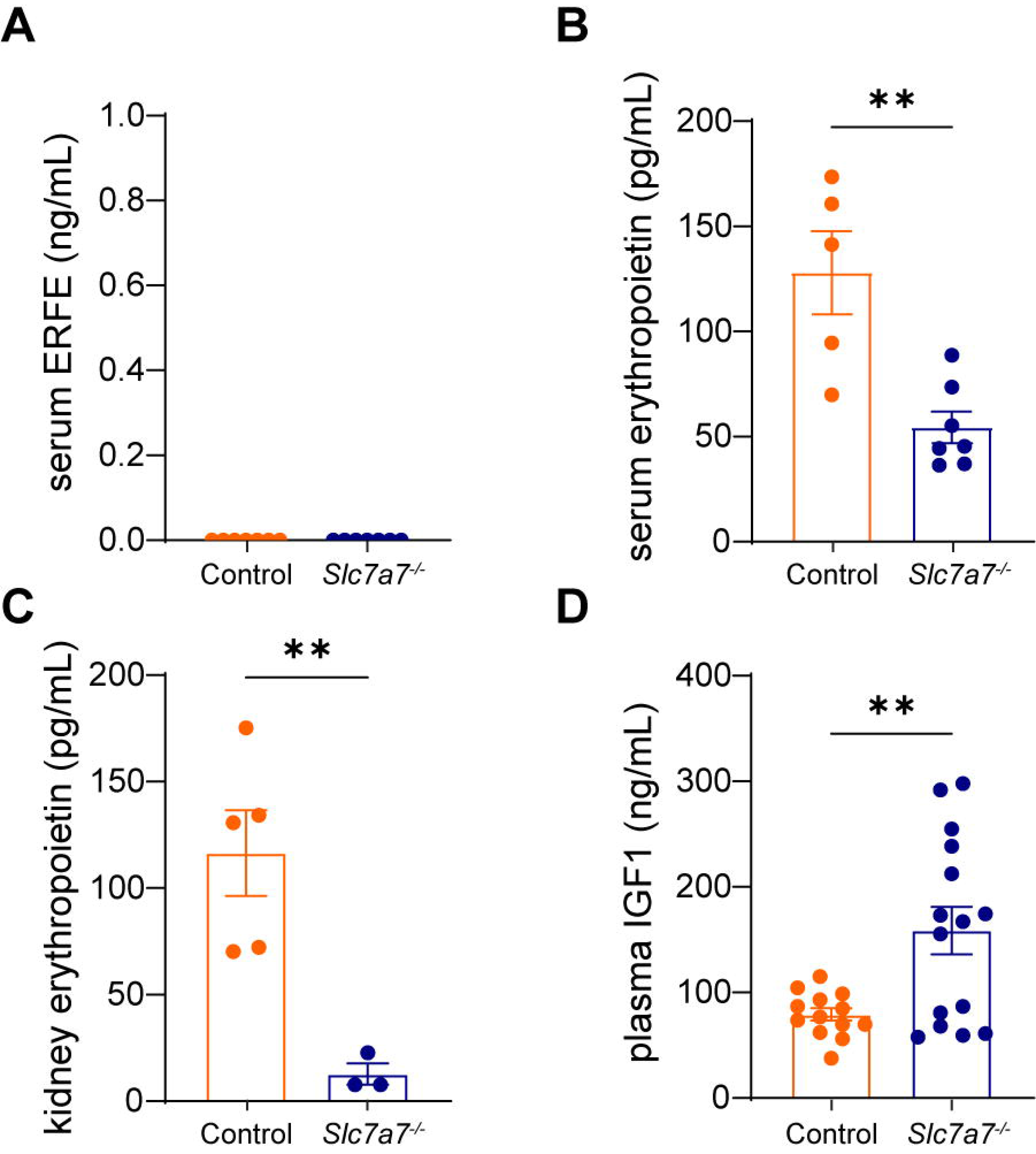
Erythropoietin levels are reduced in *Slc7a7^-/-^* mice. (A) ELISA of serum erythroferrone in control and *Slc7a7^-/-^* mice. (B) ELISA of serum erythropoietin concentration. (C) ELISA of erythropoietin concentration in total kidney extract. (D) ELISA of plasma Insulin growth factor 1. Data information: Data are mean ± SEM. **p < 0.01 (two-tailed unpaired student-t test). Each data point represents a single animal.

### Erythropoietin supplementation restores normal erythropoiesis and iron overload

To study whether the reduced systemic erythropoietin was the main cause of the hematological abnormalities and iron overload observed in *Slc7a7^-/-^* mice, we administered recombinant human erythropoietin (500 IU/kg) to both groups of mice over three consecutive days (Fig 5A) and measured erythroblast precursors, whole blood count and iron content. In agreement with a previous study (30), the administration of erythropoietin increased spleen weight by two-fold in both *Slc7a7^-/-^* and wild-type mice (Figs 5B-C and Fig S6A). However, body weight was unchanged in these two genotypes (Fig S6B). We next tested whether erythropoietin administration affects RPM number, as erythropoietin administration is known to increase the splenic macrophage population. This analysis revealed that, whereas erythropoietin increased the RPM population in control mice, it had no effect on this parameter in *Slc7a7^-/-^* mice (Fig 5D and S6C). We analyzed erythroblast precursors to determine whether erythropoietin was the cause of erythropoiesis failure in *Slc7a7^-/-^* mice (Fig 3B). All erythroblast precursors were fully recovered upon exogenous erythropoietin administration (Fig 5E), and whole blood count analysis showed that MCV and other hemogram features were comparable between control and *Slc7a7^-/-^* mice (Table 2). These findings suggest that reduced systemic erythropoietin underlies the hematological alterations of the mouse model of LPI. Interestingly, the iron phenotype was also fully reversed after erythropoietin treatment (Figs S6D-H).

**Figure 5.**
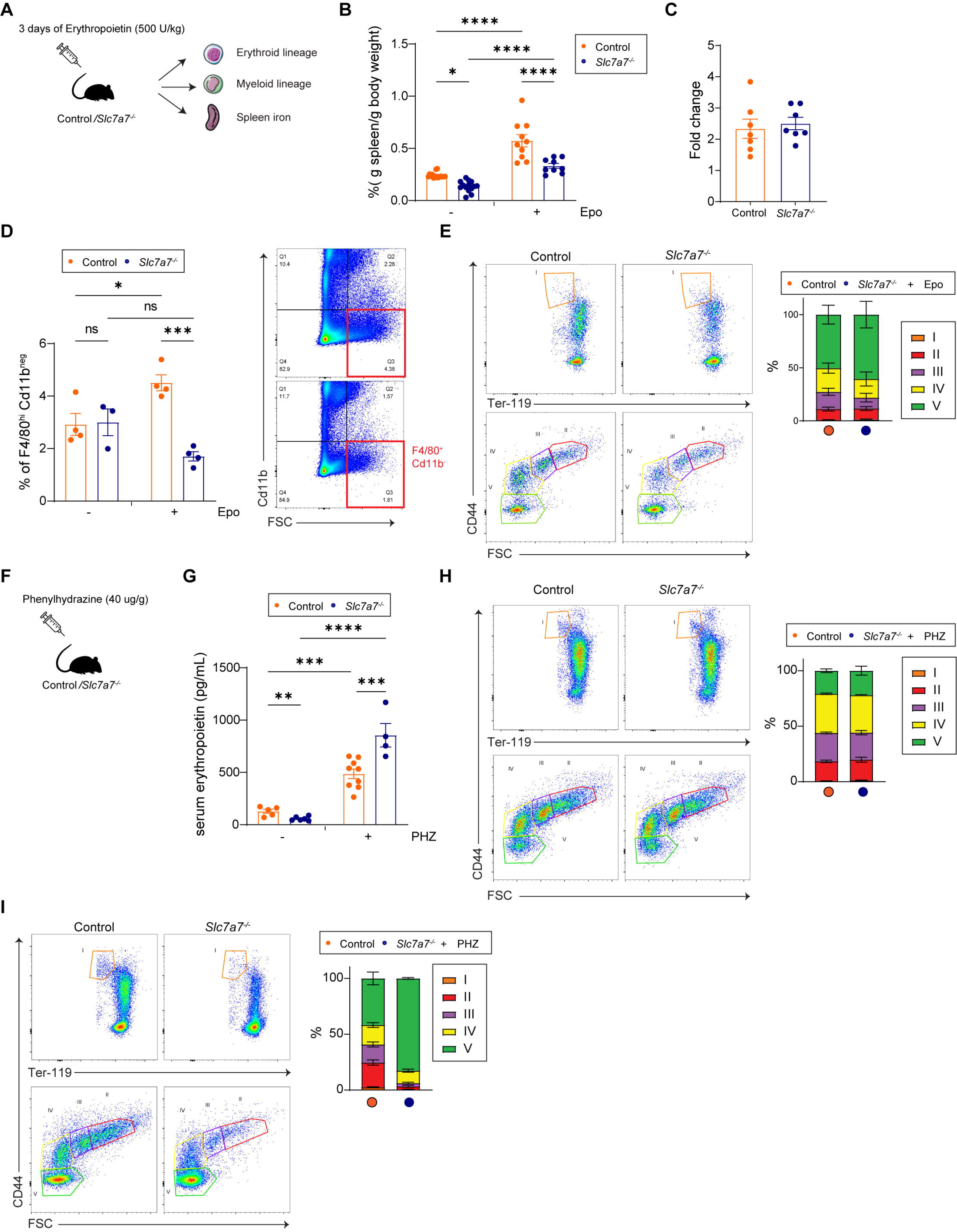
Erythropoietin is required for normal erythropoiesis and normal iron homeostasis in *Slc7a7* knockout mice. (A) Graphical scheme of erythropoietin administration in control mice and in the inducible *Slc7a7* mouse model. (B) Relative spleen weight of control and *Slc7a7^-/-^* mice with or without erythropoietin administration (*p < 0.05, ****p < 0.0001 one-way ANOVA). (C) Fold change of spleen weight increase after erythropoietin administration in control and *Slc7a7^-/-^* mice. (D) Flow cytometry analysis of red pulp macrophage (F4/80^+^Cd11b^-^). Window in Q3 represents the red pulp macrophages population. (*p < 0.05, ***p < 0.001 one-way ANOVA). (E) Left: Representative dot plots show the gating strategy for erythroid progenitors (V, IV, III, II, and I) (Chen) from the indicated genotype after erythropoietin administration. Briefly, cells were first gated in Ter-119^+^ and further separated by CD44 vs. Forward Scatter (FSC-A). Right: Percentage of the cell populations analyzed. (F) Graphical scheme of erythropoietin administration in control mice and in the inducible *Slc7a7* mouse model. (G) ELISA of serum erythropoietin concentration (**p < 0.01, ***p< 0.001, ****p < 0.0001 one-way ANOVA) (H) Left: Representative dot plots show the gating strategy for erythroid progenitors (V, IV, III, II, and I) in BM (Chen) from the indicated genotype after phenylhydrazine administration. Right: Percentage of the cell populations analyzed. (I) Left: Representative dot plots show the gating strategy for erythroid progenitors (V, IV, III, II, and I) in spleen (Chen) from the indicated genotype after phenylhydrazine administration. Right: Percentage of the cell populations analyzed. Data information: Data are mean ± SEM. *p < 0.05, **p < 0.01, ***p < 0.001, ****p < 0.0001 (two-tailed unpaired student-t test). Each data point represents a single animal.

**Table 2.** Whole blood count analysis in Slc7a7^-/-^ mice after three consecutive days of recombinant erythropoietin administration. Hemoglobin (Hb), hematocrit (Hct), mean corpuscular hemoglobin (MCH), mean corpuscular volume (MCV), and red blood cell counts (RBC) from control and Slc7a7^-/-^ mice. Data are ± SEM. NS stands for non-significant. P-val were calculated using a two-tailed Student’s t-test.

| Genotype | Hb, g/dL | HCT, % | MCH, pg | MCV, fL | RBC, 10<sup>6</sup>/μL |
| --- | --- | --- | --- | --- | --- |
| Control | 13.62 ± 2.36 | 41.35 ± 6.59 | 15.12 ± 0.42 | 46.02 ± 1.07 | 8.99 ± 1.43 |
| *Slc7a7*<sup>-/-</sup> | 15 ± 0.71 NS | 45.86 ± 1.3 NS | 14.66 ± 0.09 NS | 44.76 ± 0.73 NS | 10.26 ± 0.44 NS |

To test whether *Slc7a7^-/-^* mice synthesize erythropoietin, we induced acute hemolytic anemia in both control and *Slc7a7^-/-^* mice through a single intraperitoneal injection of phenylhydrazine (PHZ) at a dose of 40 μg/g of body weight. Our results showed that *Slc7a7^-/-^* mice had a significant increase in plasm^μ^a erythropoietin levels compared to the controls (Fig 5F-Interestingly, BM erythropoiesis (Fig 5H), FPN1 expression, serum iron and transferrin saturation were restored in the *Slc7a7^-/-^* mice (Figs S6I-J and S6M)); however, plasma ferritin and tissue iron in spleen were not recovered (Figs S6K-L) and EMH in the spleen was not triggered in the knockout mice (Fig 5I). As monocyte migration to the spleen is crucial for the generation of the stress erythropoiesis niche (31), we also quantified the number of circulating monocytes and observed it to be significantly higher in *Slc7a7^-/-^* mice after PHZ administration (Fig S6N). Given that as *Slc7a7^-/-^* knockout mice respond to hemolytic anemia but are unable to trigger EMH, we suggest that the failure of erythropoiesis and iron overload in these mice is primarily due to the lack of erythropoietin.

### *Slc7a7* knockout mice present renal dysfunction

Erythropoietin in adult mice is largely produced and secreted by the kidney (27). The cells responsible for erythropoietin production are called renal erythropoietin-producing cells (REPCs) and they are located mainly in the renal cortico-medullary junctions (32). Since *Slc7a7* is located primarily in the cortex, specifically in the epithelial cells of the proximal tubule (Fig S7A), we initially suspected that REPCs might express *Slc7a7*. To test this notion, we used tissue *in-situ* single cell RNAseq by fluor-probe hybridization and confocal microscopy (33) to perform co- localization studies of REPCs and *Slc7a7*. In steady-state conditions, *Epo* was poorly expressed in these cells (Fig S7B). Hence, to boost erythropoietin production, we challenged the mice with acute PHZ administration to induce hemolytic anemia. To assess REPCs, we analyzed the co-expression of *Epo^+^* gene with *Pdgfr +*, an interstitial fibroblast marker (34). Surprisingly, we found that cells co-expressing both genes^β^also express *Slc7a7* (Fig S7C, right) and were located around tubular cells. We also examined the expression of *Aqp2*, a protein that forms channels allowing the traffic of water molecules across the cell membrane. Staining of *Aqp2* was observed in collecting ducts and in REPCs (Fig S7B-C).

The transformation of REPCs to myofibroblasts because of fibrosis has been reported to be the primary cause of reduced erythropoietin production, a common feature of anemia caused by chronic kidney disease (35). Strikingly, end-stage renal disease has a high prevalence in patients with LPI. Therefore, we tested whether *Slc7a7^-/-^* mice developed renal and/or kidney fibrosis. Plasma analysis showed that *Slc7a7^-/-^* mice had elevated blood urea nitrogen (Fig 6A) compared to control wild-types. Furthermore, we previously published that *Slc7a7^-/-^* mice had a reduced renal function as reflected by a decreased glomerular filtration rate and diminished creatinine secretion in urine (13). We then performed histological image analysis and quantification of Sirius Red staining, which is used to detect collagen networks. A tendency of increased collagen network was observed in interstitial cells in the kidneys from *Slc7a7^-/-^* mice (Fig 6B). Mitochondrial dysfunction is strongly correlated with acute kidney injury, chronic kidney disease, and diabetic nephropathy (36). To test whether the kidneys of the knockout mice had reduced mitochondrial respiration we performed high respirometry studies using complex-specific inhibitors and substrates. We found that mitochondrial respiration was significantly reduced in the cortex from *Slc7a7^-/-^* mice (Fig 6C) in the absence of changes in mitochondrial mass, assessed by mtDNA copy number (Fig 6D). Overall, these data suggest that *Slc7a7* is dispensable for erythropoietin production and that in steady-state conditions, renal dysfunction, is the underlying cause of reduced erythropoietin production.

**Figure 6.**
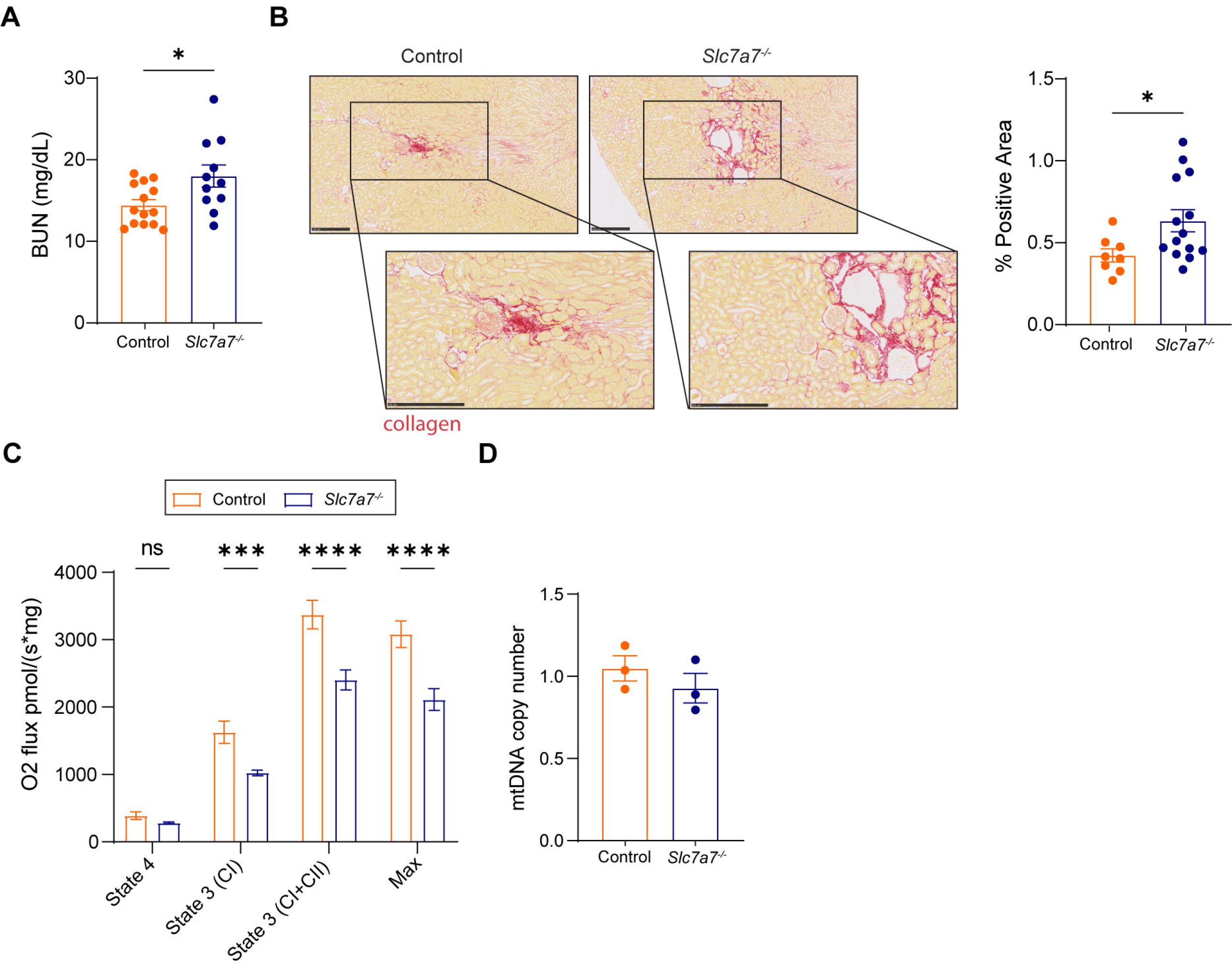
*Slc7a7^-/-^* mice develop kidney damage and fibrosis. (A) Blood urea nitrogen (BUN) concentrations in control and *Slc7a7^-/-^* mice. (B) Left panel: Representative photographs of Sirius red staining in kidney sections from control and *Slc7a7^-/-^* mice. Right panel: Quantification of Sirius red staining. (C) Mitochondrial respiratory capacity in mitochondrial extract from the cortex of control and *Slc7a7^-/-^* mice. (D) Relative mitochondrial DNA content in *Slc7a7^-/-^* and control mice. Data information: Data are mean ± SEM. *p < 0.05, **p < 0.01, ***p < 0.001, ****p < 0.0001 (two tailed unpaired t-student test). Each data point represents a single animal.

### Citrulline treatment could mask iron overload in LPI patients

Several *Slc7a7* mutations responsible for LPI have been described in different geographic locations, being more prominent in Finland and Italy (37). Since the previously described LPI mouse model recapitulates the metabolic deficiencies of the disease (13), we addressed whether different mutations from an Italian cohort are associated with high serum iron and low erythropoietin serum levels. To this end, we evaluated serum samples from 10 patients with LPI between 9 and 55 years of age, both males and females. Ten mutations were identified in this cohort, where 9 individuals carried homozygous mutations (70%) and 3 individuals were double heterozygous (30%). A summary of patients’ mutations is provided in Supplementary Table S1. Symptomology was highly heterogeneous among patients, 3 of the patients had hyperammonemia (30%), 7 had pulmonary disease evaluated by TC alterations (70%) and 4 had renal disease (40%). LPI disease is commonly treated with a low protein diet and citrulline administration (4) and in severe cases of hyperammonemmia, ammonia scavengers are also administered. Regarding the treatment, 5 of 10 patients were on a low protein diet (50%) and citrulline therapy (90%). However, ammonia scavengers were administered to only 7 patients (70%). A summary of the patients’ symptomology can be found in Supplementary Table S1.

Patients with LPI receiving citrulline therapy did not have significantly elevated serum iron and transferrin saturation compared to the reference range (Table S2). Interestingly, the only patient not under citrulline therapy had elevated serum iron levels (155 μg/dL) compared to the rest of the treated patients, and in the higher reference range. However, statistical analysis was not feasible since the sample size was too small. We then observed that 70% of the patients had elevated serum ferritin and this increase did not correlate with the age of the patients or their gender (Table S2). These findings suggested that patients with LPI develop iron overload on uncontrolled situations such as when not receiving treatment.

Next, whole blood count analysis showed that 50% of patients with LPI had elevated RBC counts, 10% had lower RBC counts, and the remaining 40% had normal levels as compared to the reference values (Table S2). We observed that MCV was reduced only in 20% of the patients, a trait significantly diminished in our mouse model (Table 1). The difference between the human disease and the mouse model could be explained by the fact that our mouse model was not treated with citrulline, which could have an impact in iron and whole blood count values.

Some patients with LPI reported cytopenia (reduced number of blood cells). We therefore studied whether this Italian cohort presented cytopenia and found that 2 individuals had diminished white blood counts (20%). In addition, neutrophils and lymphocytes were reduced in 2 patients (20%), whereas monocytes were increased in 2 (20%). A critical indicator of an inflammatory phenotype is lactate dehydrogenase (LDH) and ferritin plasma levels. LDH levels were increased in all 10 patients (100%) and ferritin in 7 (70%). These data highlight the pleiotropic phenotype of LPI disease and indicate that immunological and hematological disorders are key in the progression of the disease.

In our mouse model, we showed that *Slc7a7* ablation causes iron overload. However, this was observed in only one patient belonging to the cohort. To test whether citrulline ameliorates iron overload in LPI mice, we treated *Slc7a7^-/-^* and control mice with citrulline and then compared iron, transferrin and ferritin and erythropoietin in serum. After citrulline therapy the iron phenotype was fully restored in the knockout mice (Figs S8A-F), thereby highlighting how *Slc7a7^-/-^* mice nicely parallel human LPI. Importantly, erythropoietin plasma levels were also normalized after citrulline administration (Fig S8G), paralleling the normal levels of erythropoietin observed in human patients following citrulline therapy (Table S2).

## Discussion

Inborn errors of metabolism (IEM) are a group of rare genetic disorders caused by mutations in metabolic enzymes or transport proteins (38). Most of these conditions are rare recessive inherited disorders, which pathophysiology are poorly understood (39). Typically, IEM are diagnosed during the first weeks after birth. Yet, in most cases, there is no definitive cure, and symptoms are alleviated through different treatments. To date, LPI is an IEM with poor prognosis and no cure. However, citrulline supplementation and ammonia scavengers can bypass the metabolic deficiencies. While several studies have reported that the metabolic condition derived from *Slc7a7* loss of function may cause immunological complications (i.e., PAP) (13,29,40), other studies disputed these observations (41). We hypothesized that loss of *Slc7a7* in renal tubular cells is the underlying cause of the hematological and immunological alterations of LPI. This hypothesis is supported by the recent clinical data on a kidney transplant in a patient with LPI, where not only did aversion of protein ameliorate but also some of the LPI symptoms (i.e., normalized ferritin and CAA plasma levels)(42).

We reported here that the absence of *Slc7a7* in mice results in iron overload, as evidenced by increased transferrin, iron and ferritin in serum, as well as higher levels of iron in the liver and spleen. Our findings indicate that this iron overload is a secondary effect of defective erythropoiesis caused by reduced serum erythropoietin, as the generation of RBC requires most of the iron and upon reduced erythrocyte production, the unused iron accumulates causing iron redistribution. Our study also shows that erythropoietin administration can restore and ameliorate the hematological abnormalities associated with LPI in mice, such as inefficient erythropoiesis and iron overload.

Although essential for CAA transport in human alveolar macrophages and monocytes, the role of SLC7A7 in the immune and hematological complications of LPI does not explain why citrulline therapy is not fully effective in patients with LPI. Previous studies using human macrophages have suggested that the absence of *Slc7a7* in these cells exacerbates an inflammatory phenotype (43). Additionally, SLC7A7 expression was found to be substantially higher in AM compared to other arginine transporters (43), thereby suggesting that the LPI pulmonary complications (i.e., PAP) might solely be a result of the absence of SLC7A7, causing a defect in surfactant catabolism. Our results, however, challenge this view and suggest that the metabolic environment caused by the absence of *Slc7a7* in kidney and intestine epithelial cells is the main factor responsible for the severe LPI complications. Based on previous studies it is possible that renal disease (14), is at the basis of reduced erythropoietin in mice. Interestingly, here we show that, like human LPI *Slc7a7* knockout mice also present traits of renal dysfunction. In contrast, human patients with LPI do not show a clear reduction of erythropoietin in serum. This finding may be attributable to citrulline therapy masking and/or improving the defect in erythropoietin synthesis.

We envision that iron overload in mice is a secondary defect caused by inefficient erythropoiesis, as RBC production is greatest iron-demanding process in the organism (44). In contrast, iron overload was not present in human patients, except for one who was not receiving citrulline therapy and whose iron levels were in the upper range, in agreement with the previously reported iron accumulation in AM (45). Accordingly, after citrulline therapy, which rescues low CAA concentration in blood and consequently the urea cycle defect, *Slc7a7* knockout mice showed normalized iron levels and serum erythropoietin, thereby revealing the profound effect of citrulline supplementation on both erythropoietin synthesis and iron metabolism. Accordingly, similar ablation levels of *Slc7a7* in RPMs from *Slc7a7^LysM-/-^* and *Slc7a7^-/-^* mice further supported the above-mentioned results, reducing the relevance of *Slc7a7* RPM expression in iron metabolism and supporting the notion that the systemic environment is a primary cause of the immune complications of LPI in mice.

The synthesis of erythropoietin occurs mainly in the peritubular interstitial cells located in the cortico-medullary junctions of the kidney. The cells devoted to erythropoietin synthesis, known as REPCs, express pericyte markers such as PDGFR-β and their number is inversely proportional to the level of oxygen available. Despite their crucial role, REPCs are relatively scarce in steady-state conditions, thus hindering their study. The impact of CAA transporters, including *Slc7a7*, on REPCs has not been explored until now. Using RNA fluorescence *in situ* hybridization (RNA-FISH), our study demonstrates that REPCs express *Slc7a7* but that its expression seems to be dispensable for erythropoietin production.

There have been previous attempts to treat end-stage renal disease and anemia in patients with LPI using erythropoietin supplementation. Nevertheless, the lack of a beneficial effect could be due to the disease being too advanced at the time of erythropoietin administration (46). Citrulline treatment for LPI has also faced criticism for its potential to increase inflammation and macrophage activation through the catabolization of citrulline to arginine (47). However, the recent success of kidney transplant in an LPI patient is very promising (42) and fully agree with the recovery of erythropoiesis and iron overload in *Slc7a7* knockout mice after erythropoietin administration. Thus, our study sheds light on the underlying mechanism of kidney transplantation as a therapeutic strategy for patients with LPI.

This study highlights the systemic component as key for the erythropoiesis defect in *Slc7a7* knockout mice. However, it is also possible that cell-autonomous mechanisms in cells expressing *Slc7a7*, contribute to LPI pathology. In this regard, further studies are needed to fully understand the mechanisms underlying in the decreased spleen size in these mice, which contradicts the splenomegaly observed in patients with LPI (4) and the underlying mechanism of defective stress erythropoiesis in the spleen of *Slc7a7* knockout mice needs to be clarified. Yet, our study identifies for the first time a possible mechanism underlying the hematological complications observed in LPI patients and opens the possibility of erythropoietin administration as a new treatment to ameliorate LPI severe complications.

## Materials and Methods

### Human subjects

Human blood samples were obtained from patients affected by Lysinuric Protein Intolerance followed at the Division of Metabolic Diseases Bambino Gesu’ Children’s Hospital in Rome (Italy). The study was approved by the institutional Ethical Committee (2119_OPBG_2020), in agreement with the Declaration of Helsinki. Informed consent was obtained from patients or their parents. Blood donors are between 9 - 48 years old and they all self-reported to be healthy and tested negative for infectious disease.

### Animals

All animal work was conducted following established guidelines. The project (DARP n°9177) was favorably assessed by the Institutional Animal Care and Use Committee of the Parc Científic de Barcelona (IACUC-PCB), and the IACUC considered that the project complied with standard ethical regulations and mets the requirements of current applicable legislation (RD 53/2013 Council Directive; 2010/63/UE; Order 214/1997/GC).

*Slc7a7^loxp/loxp^* mice were generated by Eurogentec. To generate the *Slc7a7^-/-^* “*Slc7a7- deficient*” model, *Slc7a7^loxp/loxp^* animals were crossed with UBC-Cre-ERT2 mice from The Jackson Laboratory [29]. Mice were housed in groups of 2–5 per cage and were kept under a 12 h dark-light period. Food and water were supplied ad libitum. Animals were fed a standard diet (Teklad global 14% protein rodent maintenance diet) until tamoxifen induction, which consisted of a tamoxifen diet for one week. After the induction period, animals were maintained on a low-protein diet for 7–10 days prior to sacrifice. Control and *Slc7a7*-deficient littermates on a C57Bl6/J genetic background were sacrificed at 10–12 weeks of age by cervical dislocation. Tissues were dissected and flash-frozen in liquid nitrogen for RNA, protein, and iron quantification studies. For hematological and biochemical studies, blood was collected from a cardiac puncture in tubes containing either EDTA or heparin. The BM was flushed from femur and tibia bones.

Macrophage-specific and RBC-specific Slc7a7 LOF (*Slc7a7^LysM-/-^*, *Slc7a7^EpoR-/-^*) mice were generated by breeding mice with Slc7a7LoxP/LoxP mice with LysM-Cre (a gift from Dr. Nebreda’s group at IRB Barcelona, Barcelona, Spain), and EpoR-cre (a gift from Dr. Klingmuller group at Max-Planck-Institute, Freiburg, Germany), respectively. For experiments, an equal number of males and females were used.

Control and *Slc7a7*^-/-^ mice were intraperitoneally injected with recombinant human erythropoietin (500 U/kg/day; R&D Systems) daily for three consecutive days. Mice in the control group were injected with an equivalent amount of saline solution. For phenylhydrazine (PHZ) (Merck Life Science, 114715) treatment, mice were injected with 40 μg/g body weight of PHZ 48 h prior to sacrifice.

## METHODS DETAIL

### Flow cytometry and cell sorting

For the analysis of splenocytes and BM cells, crushed spleens and flushed BM samples were dissected and filtered through a 40-μm cell strainer. Erythroid cells were removed by incubation with ammonium-chloride-potassium lysis buffer prior to Fc-Blocking (anti-mouse CD16/32; ThermoFisher) for 15 min on ice. Antibodies for FACS analysis and sorting were used to stain splenic cells: APC-conjugated CD44, FITC-conjugated Ter-119, AF594-conjugated F4/80 (all antibodies from BioLegend) and APC-Cy7-conjugated Cd11b (BD Bioscience) for 30 min on ice. APC-conjugated CD44, FITC-conjugated Ter-119 and PE-CD71 (BD Pharmingen) were used for FACS analysis and BM cell sorting. Flow cytometry analysis was performed on an Aurora flow cytometer (Cytek Biosciences). Cell sorting (purity >90%) was carried out using a FACS Fusion II (BD Biosciences). For microarray analysis, spleens were prepared as described above and stained with FITC-conjugated CD106, APC-Cy7-conjugated Cd11b and PE-conjugated F4/80 (all from BioLegend) antibodies for purified RPMs. Cell doublets were excluded from all analyses and, when possible, dead cells were excluded using DAPI staining. Data analysis was carried out using FlowJoTM Software.

### Histological sample preparation and analysis

Liver, kidney, and spleen samples were fixed with neutral buffered formalin overnight at 4°C. All samples were embedded in paraffin. Paraffin-embedded tissue sections (2–3-μ air-dried and further dried at 60°C overnight.

For iron staining, paraffin-embedded tissue sections were dewaxed and stained with Iron Stain Kit to identify iron pigment using the Dako Autostainer Plus. When combining iron staining with F4/80 immunohistochemistry, iron staining was performed first.

For the Picrosirius Red staining, tissue samples were dewax and incubated with the mordant Thiosemicarbazide 99% (TSC) (Sigma, T33405) for 10 min, washed in distilled water and therefore incubated with 0.5% direct Red 80 (365548, Sigma) solution in Piric Acid Solution 1.3% (P6744-1GA, Sigma) for 90 min and finally rinsed with 1% acetic acid (Sigma, 320099) for 1 min. In all cases samples were dehydrated and mounted with Mounting Medium, Toluene-Free (CS705, Dako, Agilent) using a Dako CoverStainer.

Prior to immunohistochemistry, sections were dewaxed, and epitopes were retrieved using proteinase K for 5 min at RT for rat monoclonal anti-F4/80 (eBioscience) staining.

Quenching of endogenous peroxidase was performed with Peroxidase-Blocking Solution for 10 min at RT. Non-specific binding was blocked using 5% normal goat serum or normal donkey serum mixed with 2.5% bovine serum albumin diluted in the wash buffer, for 60 min at RT. The primary antibody dilution used was 1:2000, overnight. The secondary antibody used was a Biotin-SP (long spacer) AffiniPure Donkey Anti-Rat IgG (H+L) at 1:500 (in wash buffer) for 60 min, followed by amplification with Streptavidin-Peroxidase polymer at 1:1000. Antigen–antibody complexes were revealed with 3-3 -diaminobenzidine, with time exposure of 1 min. Sections were counterstained with hematoxylin an^′^ d mounted with Toluene-Free Mounting Medium, using a Dako CoverStainer. Specificity of staining was confirmed with a rabbit IgG, polyclonal - Isotype control or Normal Rat IgG Control. Immunofluorescence staining of y^+^LAT1 in kidney sections was performed as described previously (48).

Brightfield images were acquired with a NanoZoomer-2.0 HT C9600 digital scanner (Hamamatsu) equipped with a 20× objective. All images were visualized with NDP.view 2 U123888-01 software using a gamma correction set at 1.8 in the image control panel of the software.

### Reticulocyte counts

Fresh EDTA-anticoagulated blood samples were stained with New methylene blue dye (Sigma, 556416) following the manufacturer’s instructions. Films on glass slides were then performed for each sample. New methylene blue was used to evaluate aggregates of ribosomes and mitochondria in reticulocytes. The percentage of reticulocytes was calculated as followed:

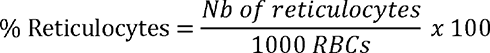

### Total kidney erythropoietin measurement

Kidneys were perfused with Hank’s balanced salts solution, and total kidney extract was obtained by mechanical digestion with PBS supplemented with protein inhibitor (Protease Inhibitor Cocktail Set III, Merck).

### Serum iron concentration

Serum iron levels were measured using a colorimetric assay (Sigma-Aldrich, MAK025) following the manufacturer’s instructions.

### Transferrin saturation measurement

Transferrin saturation was measured by a colorimetric assay that determines total iron-binding capacity (TIBC) and unsaturated iron-binding capacity (UIBC). Transferrin saturation was calculated as the percentage of serum iron divided by the TIBC (Cayman Chemical, 702230).

### Multiplex cytokine assay

Blood was extracted by cardiac puncture and then serum was prepared by centrifugation. 50 μL of serum samples was shipped to an external company (EveTechnologies) and a cytokine assay was performed. The samples were tested for the following cytokines:

### Mouse cytokine proinflammatory Focused 10-Plex discovery assay (MDF10)

GM-CSF, IFNy, IL-1β, IL-2, IL-2, IL-6, IL-12p70, MCP-1, TNF α .

### Western Blotting

Membrane proteins were extracted using a cell fractionation kit according to the manufacturer’s instructions (PromoKine, PK-CA577-K270), quantification and western blotting procedures were performed as described previously (49). GAPDH was used as a housekeeping at 1:1000 (Cell Signaling, 2118L). Proteins were detected by the enhanced chemiluminescence method (GE Healthcare Life Sciences) and quantified by scanning densitometry.

### RNA fluorescence *in situ* hybridization (RNA-FISH)

In situ 4-color RNA-FISH was performed using primer-padlock 2-oligo hybridization of the RNA targets, essentially as described (50), except that hybridization was performed in 30% formamide buffer (Merck). Briefly, ssDNA oligo probes were designed against the coding regions of target RNA transcripts (Supplementary Table S3), using Picky2.0 to identify loci without RNA secondary structure or repetitive sequence, followed by blast searching to confirm target specificity. Probe hybridization was performed overnight at 40°C with rocking in a humid chamber. Probes then underwent rolling circle amplification, polyacrylamide gel mounting, and proteinase K (Sigma-Aldrich) sample clarification before imaging. Images were acquired in a Zeiss LSM880 microscope, with processing in ImageJ using background subtraction (50 pixels) and a 2-pixel median-filter.

### Data and code availability

Microarray data have been deposited in the NCBI Gene Expression Omnibus repository under accession number GSE164827.

### Statistical analysis

Data were analyzed using GraphPad Prism Version 8 software. Statistical analysis was performed using the Student’s t-test and ANOVA, as specified in each figure legend.

## Acknowledgments

We thank Vanessa Hernández and Jorge Manuel Seco for experimental assistance and mouse handling. We also thank Prof. Ursula Klingmüller for providing us the ErGFPcre mice line. We also thank the Histopathology Core Facility, especially, Mònica Aguilera, Anais Mallen and Alicia Ferrer, for help. This work was supported by grants from the Spanish Ministry of Science and Innovation (grant SAF2015-64869-R-FEDER, RTI2018-094211-B-100 and PID2021-122802OB-100), the Ramon Areces Foundation (I.O.F.R. ARECES), and the Generalitat de Catalunya (grant 2017 SGR 961). MS acknowledges grant RTI2018-101735-B-I00 from the Spanish Ministry of Science and Innovation. RA acknowledges “Carmen de Torres” grant. The EMBO Short-Term Fellowship Program facilitated the collaboration between international groups.

---

**Supplementary Figure 1.**
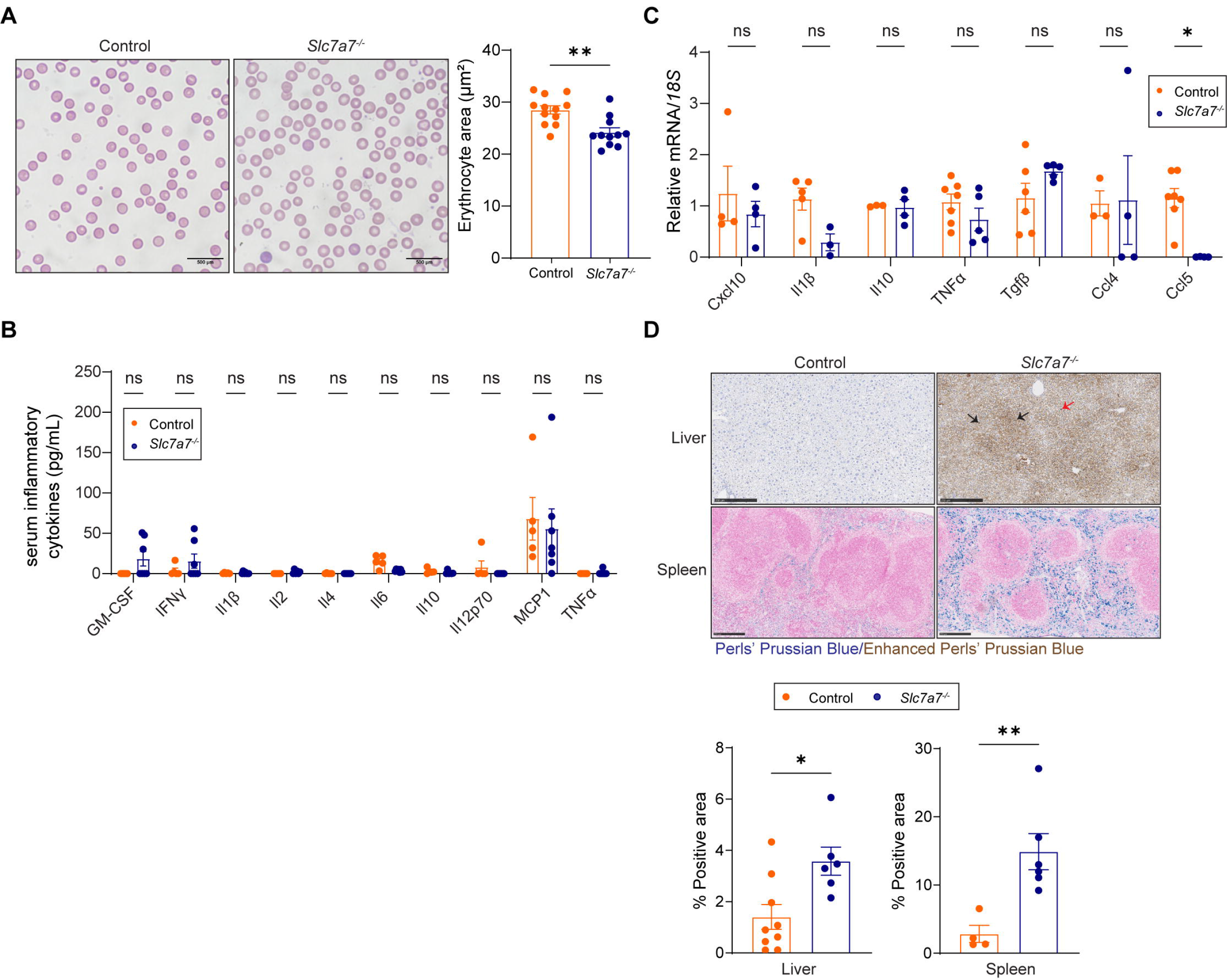

**Supplementary Figure 2.**
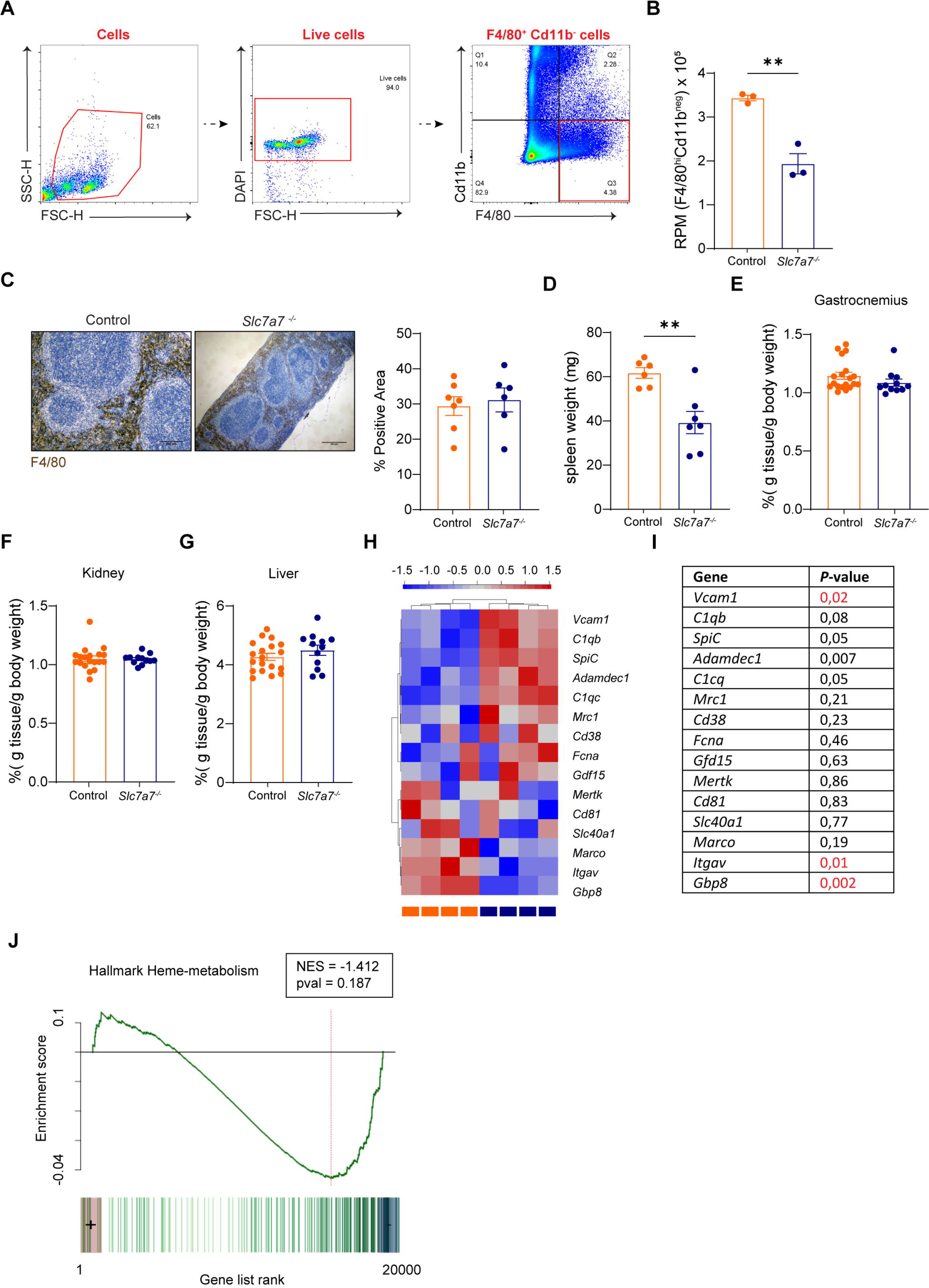

**Supplementary Figure 3.**
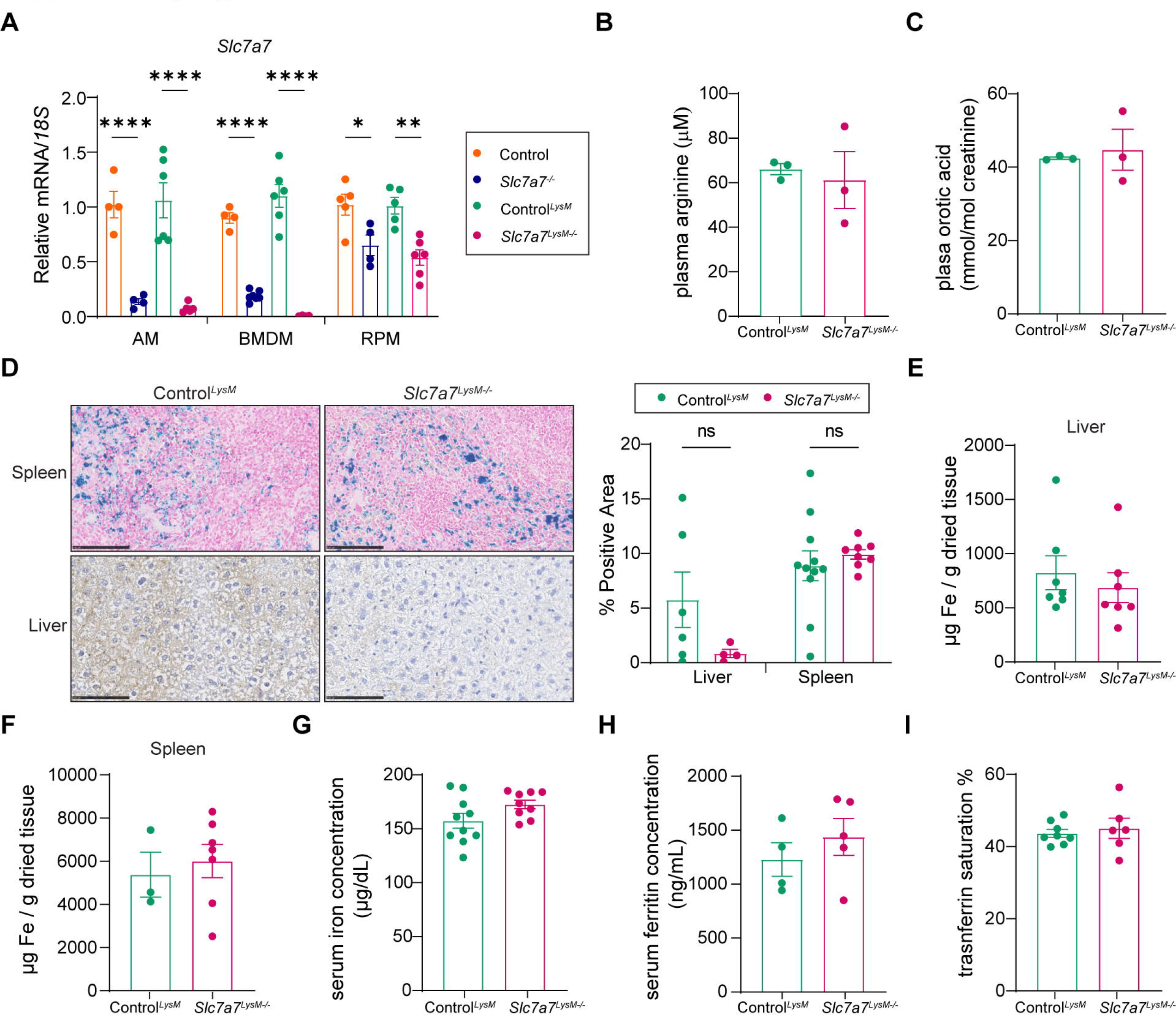

**Supplementary Figure 4.**
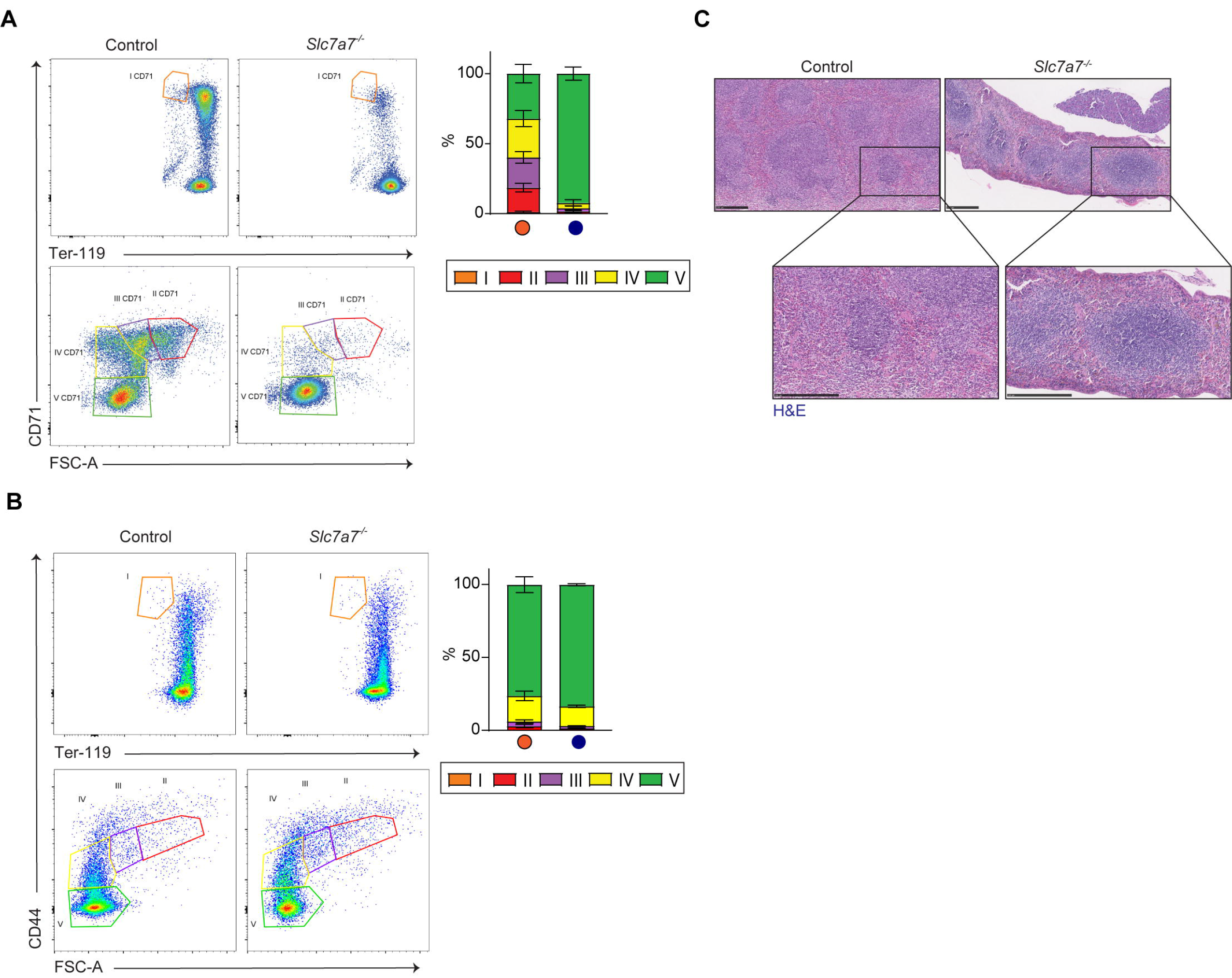

**Supplementary Figure 5.**
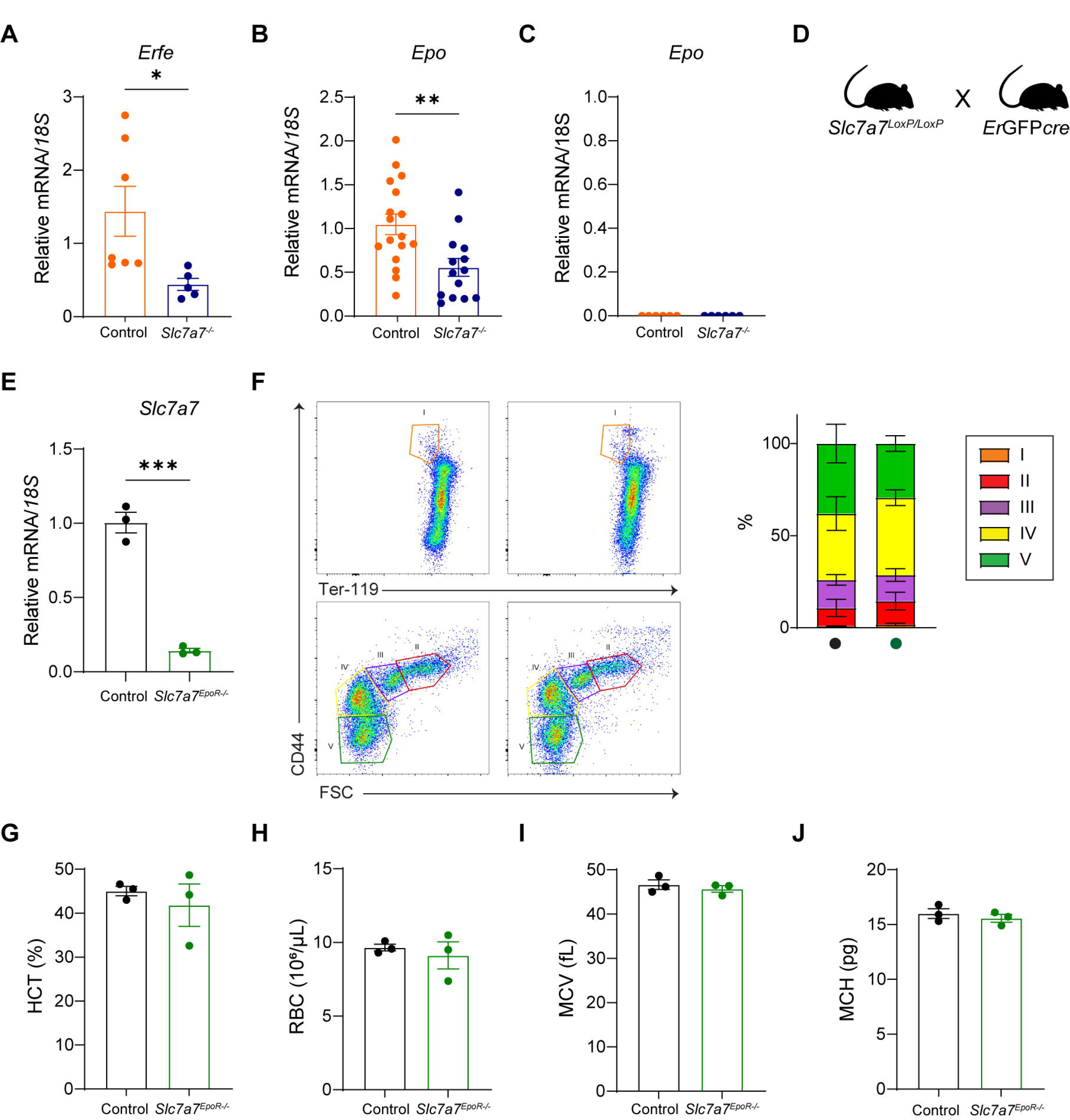

**Supplementary Figure 6.**
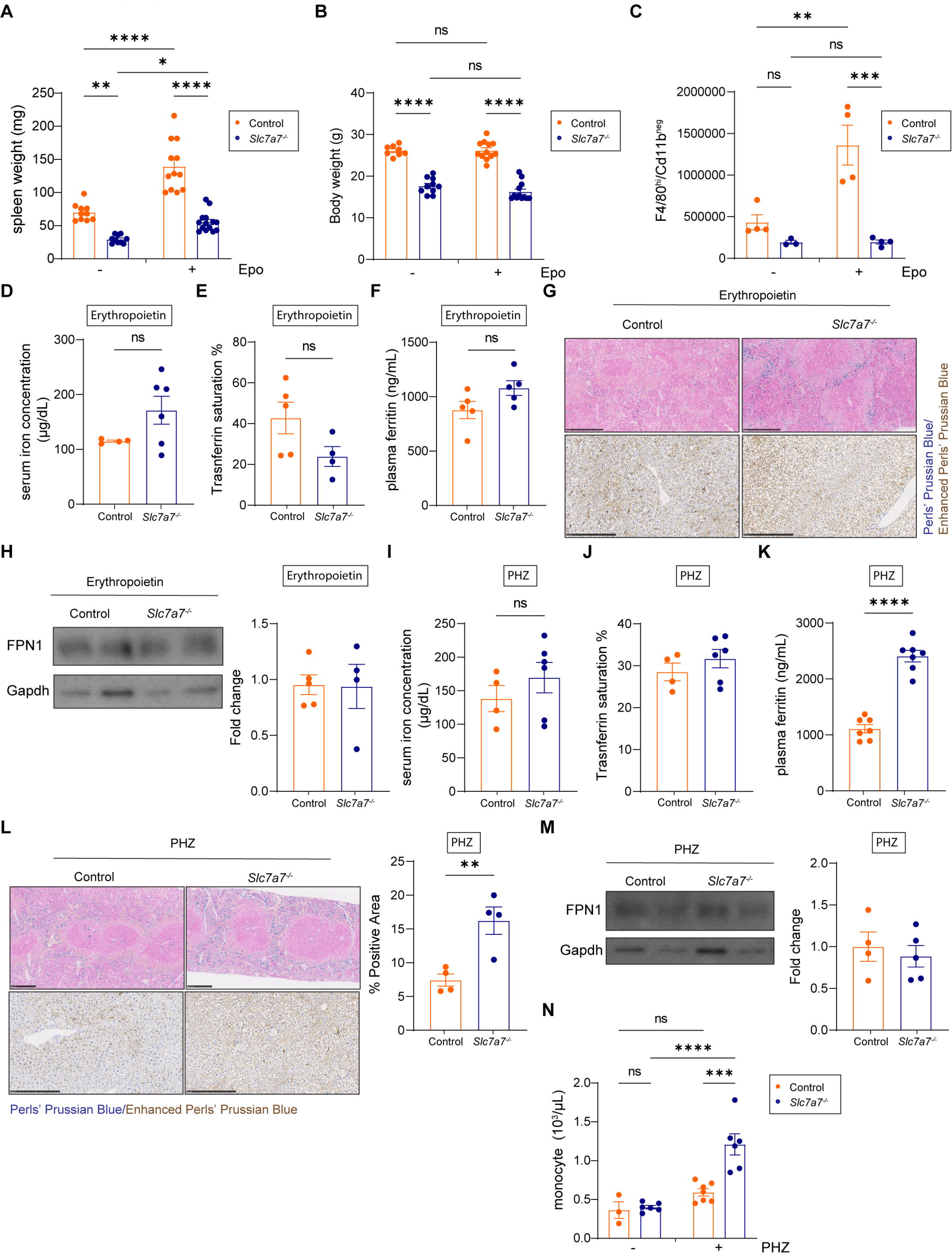

**Supplementary Figure 7.**
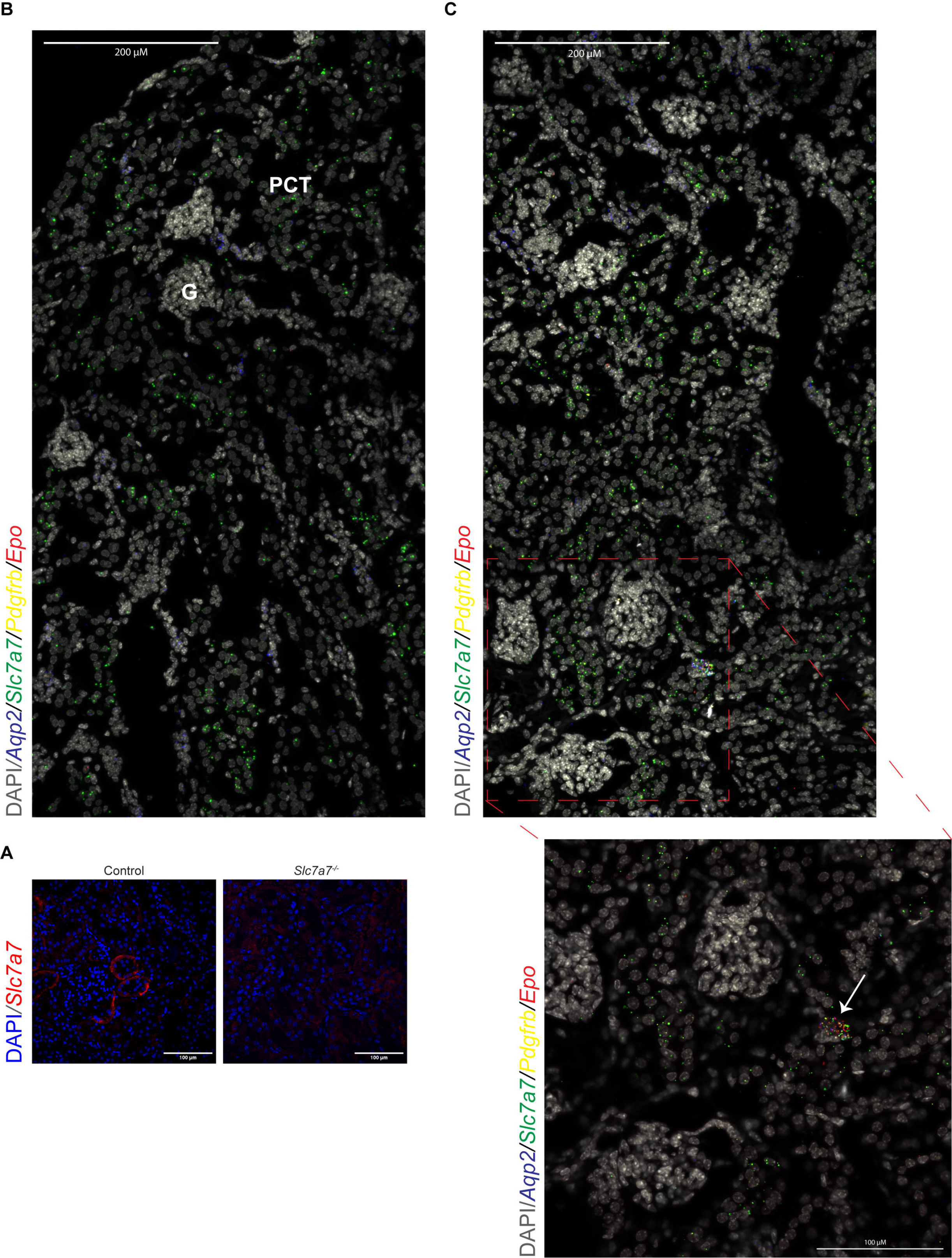

**Supplementary Figure 8.**
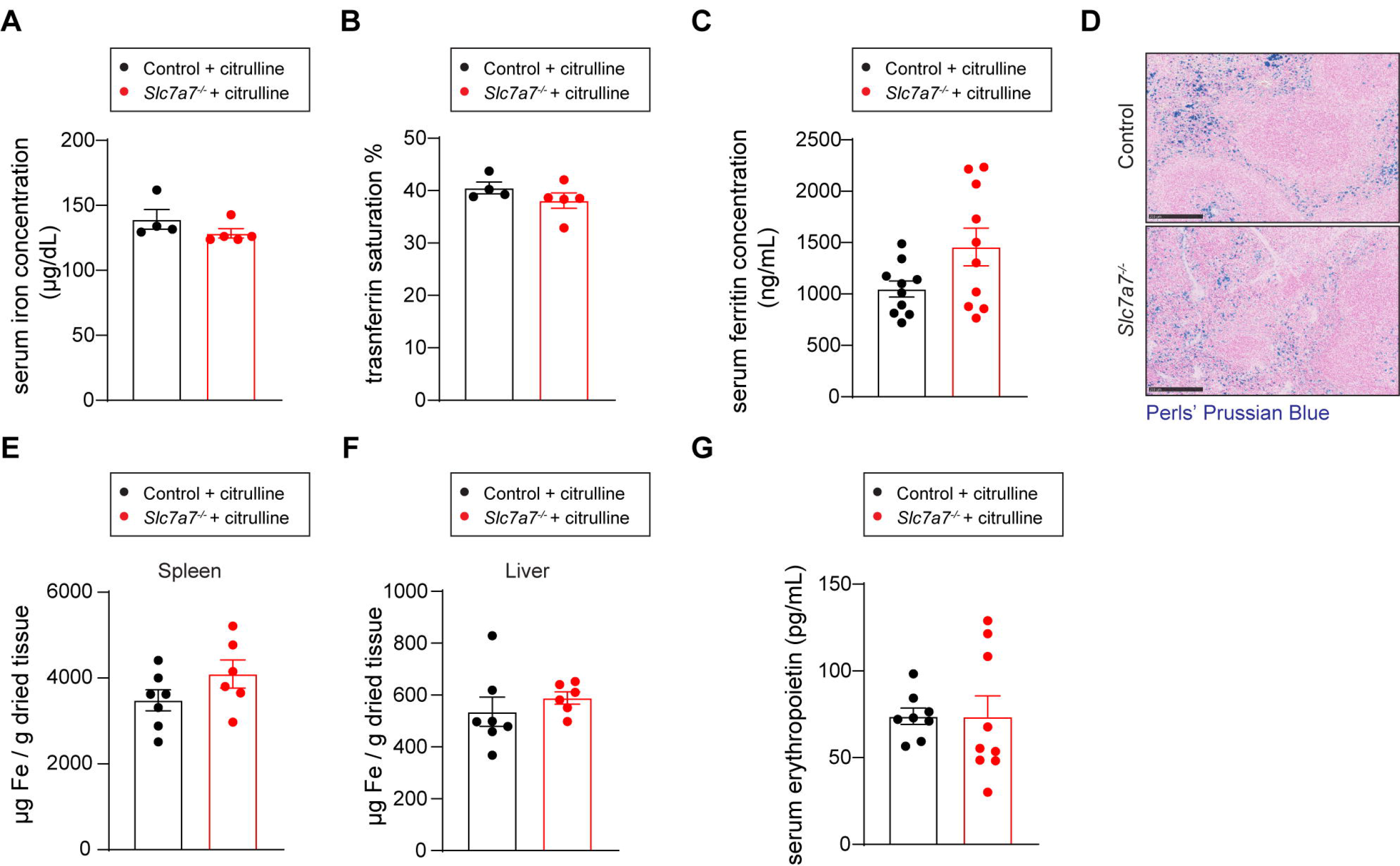

## References

1. Bröer S, Bröer A. Amino acid homeostasis and signalling in mammalian cells and organisms. Biochemical Journal [Internet]. 2017 Jun 15 [cited 2022 Mar 6];474(12):1935–63. Available from: https://pubmed.ncbi.nlm.nih.gov/28546457/

2. Palacín M, Estévez R, BERTRAN J, Zorzano A. Molecular biology of mammalian plasma membrane amino acid transporters. Physiol Rev [Internet]. 1998;78(4):969–1054. Available from: http://physrev.physiology.org/content/physrev/78/4/969.full.pdf

3. Torrents D, Mykkänen J, Pineda M, Feliubadaló L, Estévez R, Cid R de, et al. Identification of SLC7A7, encoding y+LAT-1, as the lysinuric protein intolerance gene. Nat Genet [Internet]. 1999 Mar [cited 2021 Sep 2];21(3):293–6. Available from: https://www.nature.com/articles/ng0399_293

4. Ogier de Baulny H, Schiff M, Dionisi-Vici C. Lysinuric protein intolerance (LPI): A multi organ disease by far more complex than a classic urea cycle disorder. Mol Genet Metab [Internet]. 2012 May;106(1):12–7. Available from: https://linkinghub.elsevier.com/retrieve/pii/S1096719212000352

5. Simell O. The Metabolic and Molecular Basis of Inherited Disease, Lysinuric protein intolerance and other cationic aminoacidurias. In: Scriver C. R., Beaudet A. L., Sly S. W. VD, editor. New York: McGraw-Hill, New York; 2001. p. 4933–4956.

6. Ko JM, Shin CH, Yang SW, Seong MW, Park SS, Song J. The first Korean case of lysinuric protein intolerance: presented with short stature and increased somnolence. J Korean Med Sci [Internet]. 2012 Aug;27(8):961–4. Available from: http://www.ncbi.nlm.nih.gov/pubmed/22876067

7. de Back DZ, Kostova EB, van Kraaij M, van den Berg TK, van Bruggen R. Of macrophages and red blood cells; a complex love story. Front Physiol [Internet]. 2014 [cited 2021 Jun 2];5(January). Available from: http://journal.frontiersin.org/article/10.3389/fphys.2014.00009/abstract

8. Klei TRL, Meinderts SM, van den Berg TK, van Bruggen R. From the Cradle to the Grave: The Role of Macrophages in Erythropoiesis and Erythrophagocytosis. Front Immunol [Internet]. 2017 Feb 2 [cited 2020 May 17];8(FEB):73. Available from: http://journal.frontiersin.org/article/10.3389/fimmu.2017.00073/full

9. Goodnough LT, Skikne B, Brugnara C. Erythropoietin, iron, and erythropoiesis. Blood [Internet]. 2000 Aug 1;96(3):823–33. Available from: https://ashpublications.org/blood/article/96/3/823/104348/Erythropoietin-iron-and-erythropoiesis

10. Haase VH. Hypoxic regulation of erythropoiesis and iron metabolism. American Journal of Physiology-Renal Physiology [Internet]. 2010 Jul [cited 2022 Jan 24];299(1):F1–13. Available from: https://www.physiology.org/doi/10.1152/ajprenal.00174.2010

11. Broeker KAE, Fuchs MAA, Schrankl J, Kurt B, Nolan KA, Wenger RH, et al. Different subpopulations of kidney interstitial cells produce erythropoietin and factors supporting tissue oxygenation in response to hypoxia in vivo. Kidney Int [Internet]. 2020 Oct 1 [cited 2021 Jan 25];98(4):918–31. Available from: www.kidney-international.org

12. Webster AC, Nagler E v, Morton RL, Masson P. Chronic Kidney Disease. Lancet [Internet]. 2017 Mar 25 [cited 2023 Feb 10];389(10075):1238–52. Available from: http://www.ncbi.nlm.nih.gov/pubmed/27887750

13. Bodoy S, Sotillo, Espino-Guarch, Sperandeo, Ormazabal, Zorzano, et al. Inducible Slc7a7 Knockout Mouse Model Recapitulates Lysinuric Protein Intolerance Disease. Int J Mol Sci [Internet]. 2019 Oct 24;20(21):5294. Available from: https://www.mdpi.com/1422-0067/20/21/5294

14. Parenti G, Sebastio G, Strisciuglio P, Incerti B, Pecoraro C, Terracciano L, et al. Lysinuric protein intolerance characterized by bone marrow abnormalities and severe clinical course. J Pediatr [Internet]. 1995 Feb 1 [cited 2021 Apr 9];126(2):246–51. Available from: http://www.jpeds.com/article/S002234769570552X/fulltext

15. Kell DB, Pretorius E. Serum ferritin is an important inflammatory disease marker, as it is mainly a leakage product from damaged cells. Metallomics [Internet]. 2014 [cited 2023 Feb 10];6(4):748–73. Available from: https://pubmed.ncbi.nlm.nih.gov/24549403/

16. Feng X, Scheinberg P, Wu CO, Samsel L, Nunez O, Prince C, et al. Cytokine signature profiles in acquired aplastic anemia and myelodysplastic syndromes. Haematologica [Internet]. 2011 Apr 1;96(4):602–6. Available from: http://www.haematologica.org/cgi/doi/10.3324/haematol.2010.030536

17. Fleming RE, Ponka P. Iron Overload in Human Disease. New England Journal of Medicine [Internet]. 2012 Jan 26 [cited 2023 Feb 10];366(4):348–59. Available from: http://www.nejm.org/doi/abs/10.1056/NEJMra1004967

18. McDowell LA, Kudaravalli P, Sticco KL. Iron Overload. Transfusion Medicine and Hemostasis: Clinical and Laboratory Aspects [Internet]. 2022 Apr 28 [cited 2023 Feb 10];359–60. Available from: https://www.ncbi.nlm.nih.gov/books/NBK526131/

19. Parmley RT, Spicer SS, Alvarez CJ. Ultrastructural localization of nonheme celluar iron with ferrocyanide. J Histochem Cytochem [Internet]. 1978 [cited 2023 Feb 10];26(9):729– 41. Available from: https://pubmed.ncbi.nlm.nih.gov/712049/

20. Barilli A, Rotoli BM, Visigalli R, Bussolati O, Gazzola GC, Kadija Z, et al. In Lysinuric Protein Intolerance system y+L activity is defective in monocytes and in GM-CSF-differentiated macrophages. Orphanet J Rare Dis [Internet]. 2010 Jan;5(1):32. Available from: http://www.ojrd.com/content/5/1/32

21. Haldar M, Kohyama M, So AYL, KC W, Wu X, Briseño CG, et al. Heme-Mediated SPI-C Induction Promotes Monocyte Differentiation into Iron-Recycling Macrophages. Cell [Internet]. 2014 Mar;156(6):1223–34. Available from: http://linkinghub.elsevier.com/retrieve/pii/S0092867414002761

22. Nemeth E, Tuttle MS, Powelson J, Vaughn MB, Donovan A, Ward DM, et al. Hepcidin Regulates Cellular Iron Efflux by Binding to Ferroportin and Inducing Its Internalization. Science (1979) [Internet]. 2004 Dec 17;306(5704):2090–3. Available from: http://www.ncbi.nlm.nih.gov/pubmed/15514116

23. Paulson RF, Hariharan S, Little JA. Stress erythropoiesis: definitions and models for its study. Exp Hematol. 2020 Sep 1;89:43–54.e2.

24. Tusi BK, Wolock SL, Weinreb C, Hwang Y, Hidalgo D, Zilionis R, et al. Population snapshots predict early haematopoietic and erythroid hierarchies. Nature [Internet]. 2018 Mar 21 [cited 2021 Jul 8];555(7694):54–60. Available from: http://dx.doi.org/10.1038/nature25741

25. Kautz L, Jung G, Valore E v, Rivella S, Nemeth E, Ganz T. Identification of erythroferrone as an erythroid regulator of iron metabolism. Nat Genet [Internet]. 2014 Jul 1 [cited 2021 Mar 21];46(7):678–84. Available from: http://www.nature.com/articles/ng.2996

26. Bhoopalan SV, Huang LJS, Weiss MJ. Erythropoietin regulation of red blood cell production: from bench to bedside and back. F1000Res [Internet]. 2020 [cited 2022 Jan 18];9. Available from: https://pubmed.ncbi.nlm.nih.gov/32983414/

27. Suzuki N, Obara N, Pan X, Watanabe M, Jishage KI, Minegishi N, et al. Specific contribution of the erythropoietin gene 3’ enhancer to hepatic erythropoiesis after late embryonic stages. Mol Cell Biol [Internet]. 2011 Sep 15;31(18):3896–905. Available from: http://www.ncbi.nlm.nih.gov/pubmed/21746884

28. Ratajczak J, Zhang Q, Pertusini E, Wojczyk BS, Wasik MA, Ratajczak MZ. The role of insulin (INS) and insulin-like growth factor-I (IGF-I) in regulating human erythropoiesis. Studies in vitro under serum-free conditions--comparison to other cytokines and growth factors. Leukemia [Internet]. 1998 Mar;12(3):371–81. Available from: http://www.ncbi.nlm.nih.gov/pubmed/9529132

29. Sperandeo MPM, Annunziata P, Bozzato A, Piccolo P, Maiuri L, D’Armiento M, et al. Slc7a7 disruption causes fetal growth retardation by downregulating Igf1 in the mouse model of lysinuric protein intolerance. Am J Physiol Cell Physiol [Internet]. 2007 [cited 2016 Sep 14];293(1):191–8. Available from: http://www.ncbi.nlm.nih.gov/pubmed/17376816

30. Millot S, Andrieu V, Letteron P, Lyoumi S, Hurtado-Nedelec M, Karim Z, et al. Erythropoietin stimulates spleen BMP4-dependent stress erythropoiesis and partially corrects anemia in a mouse model of generalized inflammation. Blood [Internet]. 2010 Dec 23 [cited 2022 Jan 24];116(26):6072–81. Available from: https://pubmed.ncbi.nlm.nih.gov/20844235/

31. Liao C, Prabhu KS, Paulson RF. Monocyte-derived macrophages expand the murine stress erythropoietic niche during the recovery from anemia. Blood [Internet]. 2018 Dec 13 [cited 2022 Mar 18];132(24):2580–93. Available from: /pmc/articles/PMC6293871/

32. Yamazaki S, Souma T, Hirano I, Pan X, Minegishi N, Suzuki N, et al. A mouse model of adult-onset anaemia due to erythropoietin deficiency. Nat Commun [Internet]. 2013 Oct 3 [cited 2021 Oct 15];4(1):1950. Available from: https://www.nature.com/articles/ncomms2950

33. Farack L, Itzkovitz S. Protocol for Single-Molecule Fluorescence In Situ Hybridization for Intact Pancreatic Tissue. STAR Protoc. 2020 Jun 19;1(1).

34. Urrutia AA, Afzal A, Nelson J, Davidoff O, Gross KW, Haase VH. Prolyl-4-hydroxylase 2 and 3 coregulate murine erythropoietin in brain pericytes. Blood [Internet]. 2016 Nov 24 [cited 2023 Feb 10];128(21):2550–60. Available from: https://pubmed.ncbi.nlm.nih.gov/27683416/

35. Olmos G, Muñoz-Félix JM, Mora I, Müller AG, Ruiz-Torres MP, López-Novoa JM, et al. Impaired erythropoietin synthesis in chronic kidney disease is caused by alterations in extracellular matrix composition. J Cell Mol Med [Internet]. 2018 Jan 1 [cited 2023 Feb 10];22(1):302–14. Available from: https://pubmed.ncbi.nlm.nih.gov/28857467/

36. Duann P, Lin PH. Mitochondria Damage and Kidney Disease. Adv Exp Med Biol [Internet]. 2017 May 1 [cited 2023 Feb 11];982:529–51. Available from: https://pubmed.ncbi.nlm.nih.gov/28551805/

37. Rotoli BM, Barilli A, Ingoglia F, Visigalli R, Bianchi MG, Ferrari F, et al. Analysis of LPI- causing mutations on y+LAT1 function and localization. Orphanet J Rare Dis. 2019;14(1):1–10.

38. Ferreira CR, Rahman S, Keller M, Zschocke J, Abdenur J, Ali H, et al. An international classification of inherited metabolic disorders (ICIMD). J Inherit Metab Dis [Internet]. 2021 Jan 1 [cited 2023 Feb 15];44(1):164–77. Available from: https://onlinelibrary.wiley.com/doi/full/10.1002/jimd.12348

39. Saudubray JM, Garcia-Cazorla À. Inborn Errors of Metabolism Overview: Pathophysiology, Manifestations, Evaluation, and Management. Pediatr Clin North Am [Internet]. 2018 Apr 1 [cited 2023 Feb 15];65(2):179–208. Available from: http://www.pediatric.theclinics.com/article/S0031395517301773/fulltext

40. 40. Stroup BM, Marom R, Li X, Hsu CW, Chang CY, Truong LD, et al. A Global Slc7a7 Knockout Mouse Model Demonstrates Characteristic Phenotypes of Human Lysinuric Protein Intolerance. Hum Mol Genet [Internet]. 2020 Jun 5 [cited 2020 Jun 17]; Available from: https://pubmed.ncbi.nlm.nih.gov/32504080/

41. Barilli A, Rotoli BM, Visigalli R, Bussolati O, Gazzola GC, Gatti R, et al. Impaired phagocytosis in macrophages from patients affected by lysinuric protein intolerance. Mol Genet Metab. 2012 Apr;105(4):585–9.

42. IJzermans T, van der Meijden W, Hoeks M, Huigen M, Rennings A, Nijenhuis T. Improving a Rare Metabolic Disorder Through Kidney Transplantation: A Case Report of a Patient With Lysinuric Protein Intolerance. American Journal of Kidney Diseases. 2022 Oct;

43. Rotoli BM, Barilli A, Visigalli R, Ingoglia F, Milioli M, di Lascia M, et al. Downregulation of SLC7A7 Triggers an Inflammatory Phenotype in Human Macrophages and Airway Epithelial Cells. Front Immunol [Internet]. 2018 Mar 19;9(MAR):508. Available from: http://journal.frontiersin.org/article/10.3389/fimmu.2018.00508/full

44. Kautz L, Jung G, Nemeth E, Ganz T. Erythroferrone contributes to recovery from anemia of inflammation. Blood [Internet]. 2014 Oct 16 [cited 2023 Feb 10];124(16):2569–74. Available from: https://pubmed.ncbi.nlm.nih.gov/25193872/

45. Parto K, Maki J, Pelliniemi LJ, Simell O. Abnormal pulmonary macrophages in lysinuric protein intolerance: Ultrastructural, morphometric, and x-ray microanalytic study. Arch Pathol Lab Med [Internet]. 1994 May [cited 2016 Sep 8];118(5):536–41. Available from: http://www.ncbi.nlm.nih.gov/pubmed/8192561

46. Tanner LM, Näntö-Salonen K, Niinikoski H, Jahnukainen T, Keskinen P, Saha H, et al. Nephropathy Advancing to End-Stage Renal Disease: A Novel Complication of Lysinuric Protein Intolerance. Journal of Pediatrics. 2007;150(6).

47. Olgac A, Yenicesu I, Ozgul RK, Biberoğlu G, Tümer L. Lysinuric protein intolerance: an overlooked diagnosis. Egyptian Journal of Medical Human Genetics [Internet]. 2020 Dec 1 [cited 2023 Feb 10];21(1):1–4. Available from: https://link.springer.com/articles/10.1186/s43042-020-00084-2

48. 48. Espino Guarch M, Font-Llitjós M, Murillo-Cuesta S, Errasti- Murugarren E, Celaya AM, Girotto G, et al. Mutations in L-type amino acid transporter-2 support SLC7A8 as a novel gene involved in age-related hearing loss. Elife [Internet]. 2018 Jan 22 [cited 2018 Dec 2];7:e31511. Available from: https://elifesciences.org/articles/31511

49. Nairz M, Schleicher U, Schroll A, Sonnweber T, Theurl I, Ludwiczek S, et al. Nitric oxide– mediated regulation of ferroportin-1 controls macrophage iron homeostasis and immune function in Salmonella infection. J Exp Med. 2013 May 6;210(5):855–73.

50. Wang X, Allen WE, Wright MA, Sylwestrak EL, Samusik N, Vesuna S, et al. Three-dimensional intact-tissue sequencing of single-cell transcriptional states. Science (1979). 2018 Jul 27;361(6400).

