## Supplementary data for "Defective *Slc7a7* transport reduces erythropoietin compromising erythropoiesis and iron homeostasis"

**This PDF file includes:**

Supporting materials

Figures S1 to S8

Tables S1 to S3

SI References

Supporting Information Text

**Materials and methods.**

**Colony Forming Units Assay (CFUs)**

12500 freshly sorted LSK (Lin^-^Sca^-^1^+^cKit^+^) cells were isolated from femur and tibiae from control and *Slc7a7^-/-^* and cultured in MethoCult M3434 (StemCell technology). The number and morphology of the colonies was assessed with phase contrast microscope after 8 days of culture.

**Respirometry studies**

Respirometry studies were performed in fresh isolated mitochondrial extract from kidney cortex using high-resolution respirometry (Oroboros Oxygraph-2k, Oroboros Instruments) as previously described (Sebastian et al., 2012, PNAS)

**Mitochondrial DNA content**

Genomic DNA was extracted from kidney tissue using DNeasy Blood and Tissue kit (Qiagen, 69504) following the manufacturer’s instructions. Mitochondrial and nuclear DNA content were assessed by measuring 16s, COX2 and HK2, UCP2 respectively by real-time PCR platform (Applied Biosystems) and the SYBRc® Green PCR Master Mix. The following primers were used: mouse *16s* forward 5’- CCGCAAGGGAAAGATGAAAGAC-3’ and reverse 5’- TCGTTTGGTTTCGGGGTTTC-3’, mouse *cox2* forward 5’- GTTGATAACCGAGTCGTTCTGC-3’ and reverse 5’-CCTGGGATGGCATCAGTTTT-3’, mouse *Hk2* forward 5’-TCTGGCTCTGAGATCCATCTTCA-3’ and reverse 5’-CCGGCCTCTTAACCACATTCC-3’, mouse *Ucp2* forward 5’-CTACAGATGTGGTAAAGGTCCGC-3’ and reverse 5’-GCAATGGTCTTGTAGGCTTCG-3’.

**Plasma measurements**

Commercial enzyme-linked immunosorbent assay kits were used to determine ferritin (Abcam, ab157713) and Insulin growth factor 1 (Abnova, KA0493).

**Serum measurements**

Commercial enzyme-linked immunosorbent assay kits were used to determine erythropoietin (R&D Systems, MEP00B), erythroferrone (Intrinsic LifeScience, ERF-200) and hepcidin (Intrinsic LifeScience, HMC-001) proteins in fresh serum.

**Tissue iron content**

Liver and spleen non-heme iron content was measured using the bathophenanthroline colorimetric method. Mouse tissues were dried at 45ºC for 3 days, weighed, and then digested for 48 h at 65ºC in 10% TCA/10% HCl to allow deproteinization of non-heme iron. Diluted extracts were added to 0.01% bathophenanthroline disulfonic acid, 0.1% thioglycolic, 7M sodium acetate solution, and absorbance at 535 nm was measured using an Ultrospec 3100pro spectrophotometer (Amersham Biosciences). The iron content of samples was obtained by interpolation from a standard curve and calibration to the weight of dried material.

**Cultures of primary bone marrow macrophages (BMDMs)**

BM cells from 12-week-old mice were flushed from femurs and tibiae. The cell suspension was lysed for 5 min in erythrolysis buffer (R&D Systems) at RT and then washed, resuspended, and cultured for 7 days in DMEM supplemented with 10% heat-inactivated FBS, 50 U/mL penicillin, 50 μg/mL streptomycin, and 50 ng/mL of recombinant M-CSF (Peprotech) or 30% of L-Cell conditioned medium. Six days after seeding, cells were harvested and re-seeded with the previously mentioned conditioned medium for 24 h. To deplete arginine, arginine-free media was used (DMEM for SILAC, Thermo Fisher).

**Mouse whole blood count analysis**

Blood was collected from cardiac puncture into EDTA microtubes. For erythrocyte area, fresh EDTA-anticoagulated blood samples were used for blood smears. Then, samples were stained with Diff-Quik staining. Finally, erythrocyte area was assessed with ImageJ software.

**Amino acid content**

Briefly, amino acids were determined by ion exchange chromatography with ninhydrin derivatization and spectrometric detection (Biochrom 30, Chromsystems, Cambridge, UK). Plasma (300 µL) were deproteinized with sulphosalicylic acid containing L-norleucine as internal standard (final concentration 100 µmol/L). After centrifugation, 200 µL of supernatant were adjusted to pH = 2.1 with lithium hydroxide, and then, injected onto the liquid chromatograph. Urinary orotic acid was analyzed following a spectrometric procedure (458 nm), by reacting with para-dimethylaminobenzaldehyde.

**Microarray analysis**

For gene expression analysis of RPMs, total RNA was isolated from purified cells using magnetic beads and the Agencourt RNA Clean XP Kit (Beckman Coulter). Quality and quantity were assessed using a Bioanalyzer 2100 (Agilent Technologies, Palo Alto, CA). RNA was amplified for 22 cycles and purified using the PureLink Quick PCR Purification Kit (Invitrogen) in the Genomic Facility of IRB Barcelona.

*Pre-processing of microarray data*

Microarray datasets were processed separately using the R packages affy and affyPLM from Bioconductor. Raw cell files data were processed using RMA and annotated using the information available on the Affymetrix-Thermo Fisher web page. Standard quality controls were performed to identify abnormal samples regarding: a) spatial artefacts in the hybridization process (scan images and pseudo-images from probe-level models); b) intensity dependence of differences between chips (MvA plots); c) RNA quality (RNA digest plot); d) global intensity levels (boxplot of perfect match log-intensity distributions before and after normalization and RLE plots); and e) anomalous intensity profile compared to the rest of the samples (NUSE plots, Principal Component Analysis).

*Differential expression*

Differential expression analysis was performed using a linear model with empirical shrinkage, as implemented in Limma R package. This model included the batch scanning for statistical control. Adjustment by multiple comparisons was performed using the Benjamini-Hochberg method.

*Biological enrichment analysis*

Genes quantified in the microarray study were annotated according to the Broad Hallmark gene set collection. Broad Hallmark sets were translated to mouse homologous genes using the R package biomaRt.

Functional enrichment analyses were performed using a modification of ROAST, a rotation-based approach implemented in the Limma R package, which is especially suitable for small experiments. Such modifications were implemented to accommodate the proposed statistical re-standardization in the ROAST algorithm, which enables its use for competitive testing. The MaxMean statistic was used for testing gene-set enrichment of Broad Hallmark. For each gene, the most variable probeset within each gene was used in these analyses (median absolute deviation). The results of these analyses were adjusted by multiple comparisons using the Benjamini-Hochberg False Discovery Rate method.

*Clustering and visualization*

The expression of selected genes was graphically represented in a heatmap with the heatmap R package, using a blue to red gradation, where red indicates the highest expression and blue corresponds to the lowest expression values. Firstly, the expression data were summarized to the gene level using the most variable probeset mapping to the same gene (median absolute deviation), and expression values were centered and scaled gene-wise. Genes and samples were clustered using the Ward agglomeration method and the correlation and Euclidean distances, respectively. To gain clarity in the graphic, the most extreme values were truncated to -1.5 and 1.5.

All analyses were carried out using R and Bioconductor.

**RNA extraction and quantitative real-time PCR**

Mice were killed by cervical dislocation, and tissues were immediately frozen for RNA isolation. Total RNA was isolated from purified cells using magnetic beads and the Agencourt RNA Clean XP Kit (Beckman Coulter). Quality and quantity were assessed using a Bioanalyzer 2100. RNA was amplified for 22 cycles and purified at the IRB Functional Genomic Facility using the PureLink Quick PCR Purification kit (Invitrogen). Amplification was performed using the ABI Prism 7900 HT real-time PCR platform (Applied Biosystems) and the SYBRc® Green PCR Master Mix. Gene expression levels were normalized with 18S as housekeeping gene. The primer sequences are as follows: mouse *Slc7a7* forward 5’-TCAACAGCACCAAGTATGAAGTG-3’ and reverse 5’- AGCCCAGATGACCAGTGAGA-3’ mouse *18S* forward 5’-GTAACCCGTTGAACCCCATT-3’ and reverse 5’-CCATCCAATCGGTAGTAGCG-3’, mouse *Hamp1* forward 5’-TGCAACAGATACCACACTG-3’ and reverse 5’-CCTATCTCCATCAACAGAT-3’, mouse *Epo* forward 5’-TGGTCTACGTAGCCTCACTTCACT-3’ and reverse 5’-TGGAGGCGACATCAATTCCT-3’, mouse *Cxcl10* forward 5’-CCAAGTGCTGCCGTCATTTTC-3’ and reverse 5’- GGCTCGCAGGGATGATTTCAA-3’, mouse *il1b* forward 5’-GGTCAAAGGTTTGGAAGCAG-3’ and reverse 5’-TGTGAAATGCCACCTTTTGA-3’, mouse *il10* forward 5’-TGGCCCAGAAATCAAGGAGC-3’ and reverse 5’-CAGCAGACTCAATACACACT-3’, mouse *Tnfα* forward 5’-GCTGAGCTCAAACCCTGGTA-3’ and reverse 5’-CGGACTCCGCAAAGTCTAAG-3’, mouse *Tgfβ* forward 5’-GGGCTACCATGCCAACTTCTG-3’ and reverse 5‘-GAGGGCAAGGACCTTGCTGTA-3’*,* mouse *Il12β* forward 5’-TGGTTTGCCATCGTTTTGCTG-3’ and reverse 5’-ACAGGTGAGGTTCACTGTTTCT-3’*,* mouse *Ccl4* forward 5’-TTCCTGCTGTTTCTCTTACACCT-3’ and reverse 5’-CTGTCTGCCTCTTTTGGTCAG-3’*,* mouse *Ccl5* forward 5’- GCTGCTTTGCCTACCTCTCC-3’ and reverse 5’-TCGAGTGACAAACACGACTGC-3’*, mouse Erfe(Fam132b)* forward 5’- ATGGGGCTGGAGAACAGC-3’ and reverse 5’-TGGCATTGTCCAAGAAGACA-3’.

**Fig. S1. Erythrocyte area, serum cytokine expression and Pearls’ Prussian blue in liver and spleen section related to Figure 1**. (A) Erythrocyte area (μm^2^) calculated in blood smear preparation. Scale bars: 500 μm. (B) Serum from control and *Slc7a7^-/-^* mice were analyzed by multiplex protein assay for 10 cytokines. (C) Real-time RT-PCR analysis of sorted red pulp macrophages expression of cytokines and interleukins. (D) Upper panel: Enhanced Pearls’ Prussian blue staining in liver sections. Brown color represents iron staining in hepatocytes (black arrow) and Kupffer cells (red arrow). Middle panel: Pearls’ Prussian blue staining in spleen sections. Hematoxylin and eosin staining was used as a background. Scale bars: 250 μm. Lower panel: quantification of percentage positive area from liver and spleen sections. Data information: Data are mean ± SEM. *p < 0.05, **p < 0.01 (two tailed unpaired t-student test). Each data point represents a single animal.

**Fig. S2.** **Additional relative organ weights and RPM phenotype related to Figure 2.** (A) Gating strategy of RPM. (B) Absolut RPM quantification in control and *Slc7a7^-/-^* mice. (C) Left: Staining of the macrophage marker F4/80(brown) in spleen sections. Hematoxylin and eosin staining was used as a background. Scale bars: 200 μm. Right: quantification. (D) Absolut spleen weight of control and *Slc7a7^-/-^* mice. (E, F and G) Relative gastrocnemius, kidney and liver weight of control and *Slc7a7^-/-^* mice, respectively. (H) Heatmap from sorted RPMs showing relative expression of key differentially-expressed RPM-associated genes (CITA HALDAR). Orange boxes stands for control animals. Blue boxes represent *Slc7a7^-/-^* mice. (I) P value of the represented genes in the heatmap from Figure 2. (J) Gene set enrichment analysis (GSEA) of Heme Metabolism (Broad Hallmarks) genes in RPMs from *Slc7a7^-/-^* as compared to control mice. Data information: Data are mean ± SEM. **p < 0.01 (two tailed unpaired t-student test). Each data point represents a single animal.

**Fig. S3. Iron phenotype in macrophage-specific *Slc7a7^LysM-/-^* mice.** (A) Real-time RT-PCR analysis of *Slc7a7* gene expression in sorted alveolar macrophages (AM), bone-marrow derived macrophages (BMDM) and RPMs. (B) Plasma arginine quantification. (C) Plasma orotic acid quantification. (D) Upper panel: Pearls’ Prussian blue staining in spleen sections. Hematoxylin and eosin staining was used as a background. Lower panel: Enhanced Pearls’ Prussian blue staining in liver sections. Brown color represents iron staining in hepatocytes. Scale bars: 250 μm. (E, F) Spectrophotometric analysis of total iron content in liver and spleen from control and *Slc7a7^LysM-/-^* mice, respectively. (G) Serum iron concentration in control and *Slc7a7^LysM-/-^* mice**.** (H) Serum ferritin saturation in *Slc7a7^LysM-/-^* mice and its control littermates. (I) Serum transferrin saturation in *Slc7a7^LysM-/-^* mice and its control littermates. Data information: Data are mean ± SEM. *p < 0.05, **p < 0.01, ***p < 0.001 and ****p < 0.0001 (two tailed unpaired t-student test). Each data point represents a single animal.

**Fig. S4. Analysis of BM erythroblast precursors with CD71 antigen and extramedullary hematopoiesis. (A)** BM erythropoiesis. Left: Representative dot plots show the gating strategy for erythroid progenitors (V (*p*-value < 0,0001), IV (*p*-value < 0,001), III (*p*-value < 0,01), II (*p*-value < 0,05) and I (*p*-value > 0,9)) (Chen) from the indicated genotype. Right: Percentage of the cell populations analyzed, n = 5. **(B)** Erythroblast precursors analysis in spleen. Left: Representative dot plots show the gating strategy for erythroid progenitors (V (*p*-value = 0,129), IV (*p*-value = 0,6368), III (*p*-value = 0,9909), II (*p*-value = 0,9957) and I (*p*-value > 0,9)) (Chen) from the indicated genotype. Right: Percentage of the cell populations analyzed, n = 3. **(C)** Hematoxylin and eosin staining in spleen sections from control and *Slc7a7^-/-^* mice. Note no differences in extramedullary erythropoiesis but decreased cell cellularity in red pulp in *Slc7a7^-/-^*mice. Scale bars: 250 μm. Data information: Data are mean ± SEM. Two-tailed unpaired t-student test.

**Fig. S5. Erythropoiesis analysis and whole blood count analysis in erythropoietin-receptor specific *Slc7a7* knockout mice.** (A) Real-time RT-PCR analysis of *Erfe* gene expression in sorted erythroblasts. (B, C) Real-time RT-PCR analysis kidney and liver, respectively, of *Epo* mRNA from control and *Slc7a7^-/-^* mice. (D) Graphical scheme of EpoR knockout mouse. (E) Real-time RT-PCR analysis of *Slc7a7* gene expression in sorted erythroblasts in control and *Slc7a7^EpoR-/-^* mice. (F) BM erythropoiesis in control and *Slc7a7^EpoR-/-^* mice. Left: Representative dot plots show the gating strategy for erythroid progenitors (V (*p*-value = 0,9696), IV (*p*-value > 0,999), III (*p*-value = 0,9988), II (*p*-value < 0,9998) and I (*p*-value = 0,9)) (Chen) from the indicated genotype. Right: Percentage of the cell populations analyzed, n = 7. (G, H, I and J) HCT, RBC, MCV and MCH, respectively, in erythropoietin-receptor specific knockout mice. Data information: Data are mean ± SEM. * p < 0.05, ** p < 0.01, *** p < 0.001 (two tailed unpaired t-student test). Each data point represents a single animal.

**Fig. S6. Iron phenotype in *Slc7a7^-/-^* mice after erythropoietin and phenylhydrazine administration, related to Figure 5.** (A) Spleen weight in control and *Slc7a7^-/-^* mice after daily injection of erythropoietin at 500 U/Kg body weight, for 3 consecutive days (Saline solution used as vehicle control) (*p < 0.05, **p<0.01, ****p < 0.0001 one-way ANOVA). (B) Body weight in control and *Slc7a7^-/-^* mice as described in A (****p < 0.0001 one-way ANOVA). (C) Absolute number of RPMs (F4/80^+^Cd11b^-^) in control and *Slc7a7^-/-^* mice as described in A (**p < 0.01, ***p < 0.001 one-way ANOVA). (D, E and F) serum iron, transferrin, and ferritin, respectively, in control and *Slc7a7^-/-^* mice as described in A. (G) Pearls’ Prussian blue staining in spleen sections. Hematoxylin and eosin staining was used as a background. Scale bars: 250 μm. (H) Left panel: Western blot analysis of Fpn-1 in total homogenate from spleen tissue in control and *Slc7a7^-/-^* mice after daily injection of erythropoietin at 500 U/Kg body weight, for 3 days. Right panel: FPN-1 quantification. (I, J, and K) serum iron, transferrin, and ferritin, respectively, in control and *Slc7a7^-/-^* mice after acute dose of PHZ at μg/g body weight. (L) Pearls’ Prussian blue staining in spleen sections in PHZ treated mice. Hematoxylin and eosin staining was used as a background. Scale bars: 250 μm. (M) Left panel: Western blot analysis of Fpn-1 in total homogenate from spleen tissue in control and *Slc7a7^-/-^* mice as described in I. Right panel: FPN-1 quantification. (N) Circulating monocytes in control and *Slc7a7^-/-^* mice treated or not with PHZ. Data information: Data are mean ± SEM. ** p < 0.01, *** p < 0.001, **** p < 0.0001 (two tailed unpaired t-student test). Each data point represents a single animal.

**Fig. S7. REP cells express *Slc7a7.*** (A) Immunofluorescence staining of y^+^LAT1 (Red) in basolateral side of proximal convoluted tubules in kidney sections from control and *Slc7a7^-/-^* mice. DAPI (blue) was used to stain nuclei. Scale bar: 100 μm. (B and C) Representative images of *Epo* mRNA (red), *Aqp2* mRNA (blue), *Pdgfrb* (yellow) and *Slc7a7* (green) by kidney sections by RNA fluorescence *in situ* hybridization in kidney sections from control mice and following PHZ administration. DAPI (grey) was used to detect nuclei. G indicates glomerulus in kidney, PCT stands for proximal convoluted tubule and the arrow outlines the REPC. Scale bar: 200 μm and 100 μm.

**Fig. S8. Citrulline supplementation restores normal iron levels.** (A, B and C) serum iron, transferrin, and ferritin, respectively, in control and *Slc7a7^-/-^* mice after 10 days of citrulline administration at 1 g/L in drinking water. (D) Pearls’ Prussian blue staining in spleen as described in A. Hematoxylin and eosin staining was used as a background. Scale bars: 250 μm. (E and F) Spectrophotometric analysis of total iron content in liver and spleen as described in A, respectively. (G) Serum erythropoietin levels as described in A. Data information: Data are mean ± SEM. Each data point represents a single animal.

Table S1. Relative information to human patients with LPI.

Table S2. Hemogram, serum erythropoietin and iron-related values from human patients with LPI.

Table S3. Primer sequences used for RNA fluorescence in situ hybridization (RNA-FISH).

**SI References**

Sample References:

1. J.-M. Neuhaus, L. Sticher, F. Meins, Jr., T. Boller, A short C-terminal sequence is necessary and sufficient for the targeting of chitinases to the plant vacuole. *Proc. Natl. Acad. Sci. U.S.A.* 88, 10362–10366 (1991).
2. E. van Sebille, M. Doblin, Data from “Drift in ocean currents impacts intergenerational microbial exposure to temperature.” Figshare. Available at [https://dx.doi.org/10.6084/m9.figshare.3178534.v2. Deposited 15 April 2016](https://dx.doi.org/10.6084/m9.figshare.3178534.v2.%20Deposited%2015%20April%202016).
3. A. V. S. Hill, “HLA associations with malaria in Africa: Some implications for MHC evolution” in Molecular Evolution of the Major Histocompatibility Complex, J. Klein, D. Klein, Eds. (Springer, 1991), pp. 403–420.
